# The spatial landscape of gene expression isoforms in tissue sections

**DOI:** 10.1101/2020.08.24.252296

**Authors:** Kevin Lebrigand, Joseph Bergenstråhle, Kim Thrane, Annelie Mollbrink, Konstantinos Meletis, Pascal Barbry, Rainer Waldmann, Joakim Lundeberg

**Author notes:** These authors contributed equally to this work (Kevin Lebrigand, Joseph Bergenstråhle, Kim Thrane).

## Abstract

*In situ* capturing technologies add tissue context to gene expression data, with the potential of providing a greater understanding of complex biological systems. However, splicing variants and fulllength sequence heterogeneity cannot be characterized at spatial resolution with current transcriptome profiling methods. To that end, we introduce Spatial Isoform Transcriptomics (SiT), an explorative method for characterizing spatial isoform variation and sequence heterogeneity. We show in mouse brain how SIT can be used to profile isoform expression and sequence heterogeneity in different areas of the tissue. SiT reveals regional isoform switching of *Plp1* gene between different layers of the olfactory bulb, and use of external single cell data allowed to nominate cell types expressing each isoform. Furthermore, SiT identifies differential isoform usage for several major genes implicated in brain function (*Snap25, Bin1, Gnas*) that we independently validated by *in situ* sequencing. SiT also provides for the first time an in-depth A-to-I RNA editing map of the adult mouse brain. Data exploration can be performed through an online resource (https://www.isomics.eu), where isoform expression and RNA editing can be visualized in a spatial context.

## INTRODUCTION

Derivation of multiple transcripts via post-transcriptional modifications such as alternative splicing and RNA editing generates substantially more transcripts than there are genes. These modifications increase transcriptome complexity and have important implications for cellular function, as evidenced by their tight regulation and role in development and tissue homeostasis^1^. Alternatively spliced transcripts are particularly important in neurogenesis and brain development, contributing to the complex architecture of the mammalian central nervous system (CNS), where they regulate a vast array of neuronal functions through cell-type-specific expression patterns^2,3^. Several links have been reported between defective alternative splicing and a growing number of diseases including epilepsy, autism spectrum disorders, schizophrenia or spinal muscular atrophy^4^. Transcriptomic diversity can also be generated through adenosine-to-inosine (A-to-I) RNA editing, a process which is mediated by a specific family of enzymes called adenosine deaminases^5^. This process is involved in proper neuronal function^6^, and dysregulated and aberrant A-to-I RNA editing has also been reported in neurological and neurodegenerative diseases such as epilepsy, amyotrophic lateral sclerosis and developmental disorders^7^.

Information on the spatial distribution of post-transcriptional modifications is crucial for a better understanding of their roles in physiology and disease. Recent technological advances have enabled high-throughput quantification of gene expression in a spatial context^8^. These methods can broadly be categorized into those that detect the presence of a predefined set of target genes and those where observations stem from sampling across the entire transcriptome. The latter is required for *a priori* free exploratory analysis and novel hypothesis generation. Such methods are usually based on *in situ* capture of poly-adenylated RNA on spatially barcoded reverse transcription primers, which allows capturing all mRNAs in the transcriptome. The captured transcripts are then sequenced *ex situ*, and their spatial barcodes are used to infer their spatial origins. Even though several new such *in situ*-capture-based methods for large-scale transcriptome profiling have recently emerged^9,10,11,12^, they only assess transcripts as 3’ cDNA tags and not as complete transcripts. The fundamental reason behind this is that all those methods are based on short-read library preparation and sequencing, which inevitably implies that full-length transcript information is lost. As the largest amount of diversity in the transcriptome stems from post-transcriptional modifications, a truly comprehensive description of the transcriptome can only be obtained by characterizing the complete sequences of the transcripts.

The situation was similar in single-cell transcriptomics where until recently just the end of the cDNA was typically sequenced. Recent bioinformatics and methodological developments have enabled detection, characterization, and quantification of full-length transcripts in single-cell experiment using either short reads^13^ or long reads generated with Pacific Biosciences (PacBio)^14^ or Oxford Nanopore Technology (Nanopore)^15,16^. Long read sequencing is the only option to unambiguously access the full exonic structure of captured transcripts^17^. PacBio sequencing has a higher accuracy (>99%) than Nanopore sequencing (97% accuracy)^18^, while the latter provides a higher sequencing throughput since a single PromethION flow cell generates more than 100 million reads, whereas only 4 million usable reads are obtained with the most recent 8M PacBio SMRTcell^19^. Considering the vast amount of RNA molecules captured in current high throughput single-cell or spatial transcriptomics approaches, Nanopore sequencing is a more attractive option to generate the large number of reads required to reach the sequencing saturation needed for comprehensive transcript isoform and sequence heterogeneity exploration.

We explored whether those recent developments in single-cell transcriptomics can be applied in spatial transcriptomics to obtain spatially resolved full-length sequence information. A recent study proposed a hybrid approach in which single cell transcript isoforms were characterized by PacBio sequencing^20^. The obtained isoforms were then inferred to spatial coordinates by shallow Nanopore sequencing of spatial transcriptomics (Visium) libraries. However, the low coverage and the lack of unique molecular identifiers (UMIs) did not enable broad exploration of the spatial sequence heterogeneity of full-length isoforms nor RNA editing in the investigated tissue.

Here, we introduce Spatial Isoform Transcriptomics (SiT), a method for comprehensive spatial profiling of full-length transcripts, based on commercially available spatially barcoded *in situ*-capture arrays and Nanopore long-read sequencing. We demonstrate the workflow for two different areas of the mouse brain and show that deep long-read sequencing identifies multiple genes that display spatially distinct alternative isoform expression. We also explored full-length sequence heterogeneity and provide for the first time a global map of A-to-I RNA editing of the adult mouse brain.

## DESIGN

Currently there is no approach available to generate comprehensive spatially resolved databases of full-length RNA sequences, and define the spatial landscape of splicing and single nucleotide variations (SNVs). We introduce here Spatial isoform Transcriptomics (SiT), which combines an existing approach that yields spatial gene expression information after short read sequencing of cDNA reverse transcribed *in situ* on tissue sections with long read sequencing to generate spatially resolved full-length mRNA sequence data. We opted for Nanopore long read sequencing because the Oxford Nanopore Promethion flow cell generates the highest number of reads and this technology is therefore more cost-effective than others (I.e., PacBio sequencing). The approach is however easily transferrable to other sequencing technologies. To analyze the sequencing data, we took advantage of the fact that spatially barcoded cDNA has a similar design as single-cell cDNA with the cell barcode replaced by a spatial barcode. This similarity allowed us to adapt a strategy recently developed for the analysis of long read single cell RNA sequencing data (ScNaUmi-seq^15^) for the analysis of spatially resolved long read sequencing data. SIT enables spatial exploration of isoform expression and RNA sequence heterogeneity in an un-targeted manner, by interrogating all captured isoforms and SNVs rather than a single isoform or SNV at a time.

## RESULTS

### Spatial isoform detection enabled through *in situ* capture and long-read sequencing

We fixed fresh-frozen tissue samples on spatially barcoded glass slides using methanol. After staining and imaging, mRNA molecules were captured *in situ* and tagged with barcodes and UMIs (10xGenomics Visium). Full-length cDNA libraries were then split for preparation of 3’ short-read sequencing libraries as well as long-read Nanopore libraries (Fig.1a). Resulting gene expression data were clustered to define distinct anatomical regions used as landmarks for analysis of regional isoform usage analysis.

**Figure 1.**
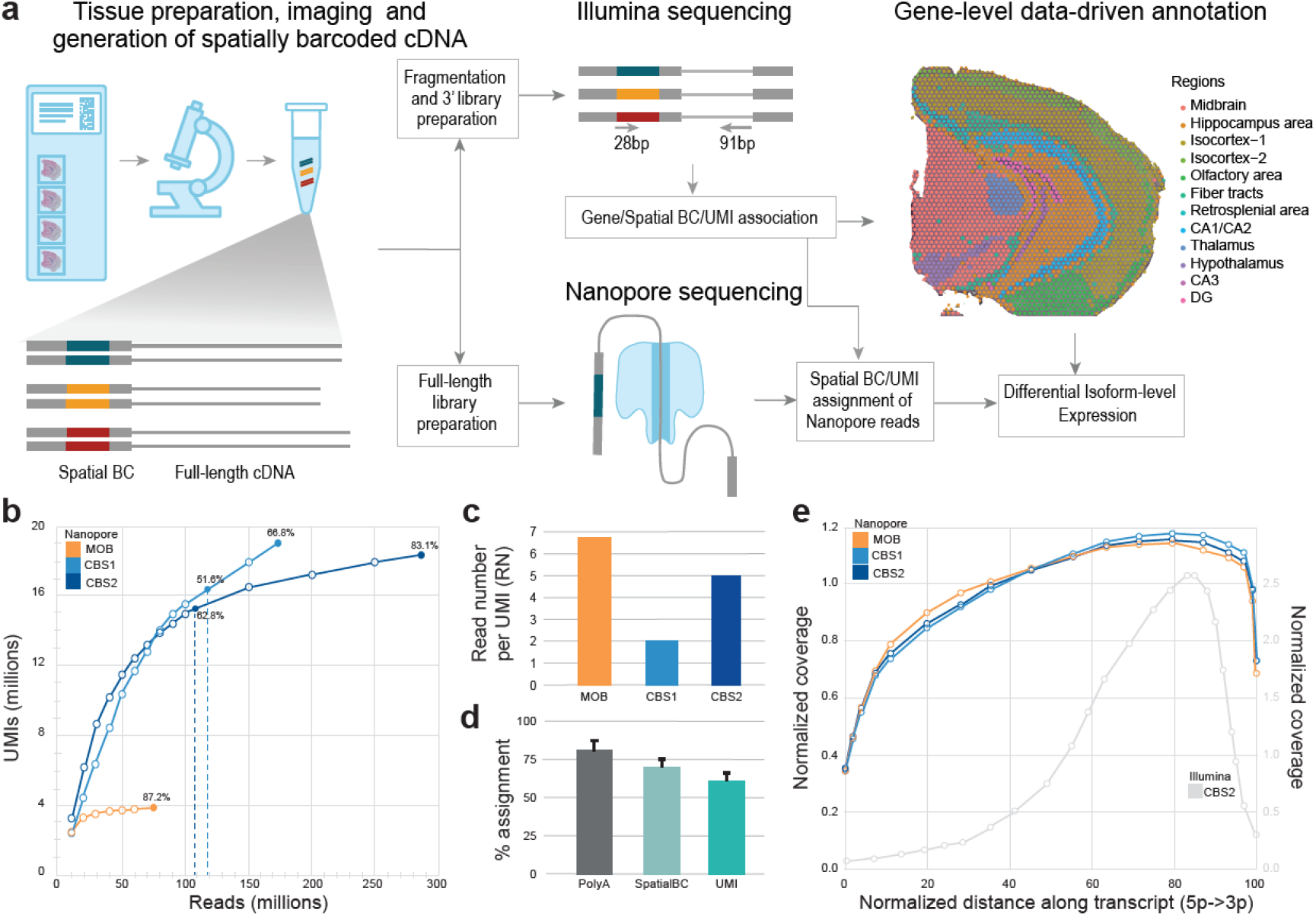
SiT methodology and datasets. **(a)** Experimental and computational steps for SiT analysis. Right side shows unsupervised gene expression clustering in a mouse coronal brain section (CBS2). **(b)** Nanopore sequencing saturation curves for three Visium samples showing the number of UMIs observed as a function of the number of Nanopore reads. Labels indicate sequencing saturations obtained with all flow cells (CBS1, CBS2, MOB) and with just one latest generation Promethion flow cell per sample (vertical dotted lines, CBS1, CBS2); **(c)** Mean read number (RN) per molecule (UMI) for each sample. **(d)** Percentage of assignment at each step of the workflow: PolyA expressed as percentage of total reads; SpatialBC expressed as percentage of PolyA found reads; UMI expressed as percentage of reads with SpatialBC. Details about the spatialBC/UMI assignment strategy are in ^15^. **(e)** Normalized transcript coverage plot for Nanopore (MOB, CBS1, CBS2) and for Illumina (CBS2) sequencing.

We demonstrate the isoform landscape *in situ* in two regions of the mouse brain: the olfactory bulb (MOB) and two coronal sections of the left hemisphere (CBS1, CBS2). We provide a dataset of 13 Nanopore PromethION flow cells with a total of 535 million reads, reaching a high long-read sequencing saturation for the three samples (Fig.1b-c). The initial Nanopore PromethION flow cells generated a median of 40 million reads, while the most recent flow cells yielded more than 100 million reads (Fig.1b, Supplementary Table 1), a throughput that provided a high sequencing saturation of 51.6% and 62.8% for CBS1 and CBS2, respectively, with just one sequencing run. We used short-read data for assignment of spatial barcodes and UMIs to Nanopore reads using the previously described scNaUMI-seq protocol^15^ (Fig.1d). Our experimental approach, that included a cDNA size selection step for full-length cDNA enrichment (Supplementary Fig.1) provided a nearly uniform representation of full-length transcripts, enabling the exploration of splicing and full-length sequence heterogeneity (Fig.1e).

### Regional isoform switching in the olfactory bulb

To investigate the spatial isoform landscape in MOB, we followed the SiT workflow depicted in Fig.1a on a fresh-frozen tissue section. We generated 253 million Illumina short reads and 74 million long reads from two PromethION flow cells reaching a sequencing saturation of 87.2% (93.1% for shortreads). For the long-read data, we applied a stringent filter to only retain molecules (UMIs) that contain all exon-exon junctions (mean exon number 6.7) of the reference isoform (Mouse Gencode vM24). Following this strategy, we unambiguously defined the full transcript structure for 2.19 million UMIs, sequenced with 59.9% of the 25.5 million spatialBC-UMI associated reads (mean 6.8 reads per UMI).

Across the tissue section, we observed 23,560 different Gencode reference isoforms of 13,291 distinct genes. Between short and long-reads, we computed a gene-level Pearson correlation of 0.91 (Supplementary Fig.2). Per spatially barcoded spot (55μm diameter), we observed a median of 1,917 UMIs corresponding to a median of 974 distinct isoforms (Supplementary Fig.3). Standard clustering of the short-read data defined five anatomic regions, as previously demonstrated^21^ (Fig.2a). Based on this unsupervised clustering, we mined for genes showing a differential isoform usage between regions and identified 36 such genes, out of which *Myl6 and Plp1* showed the most prominent patterns (Methods, Supplementary Table 4).

**Figure 2.**
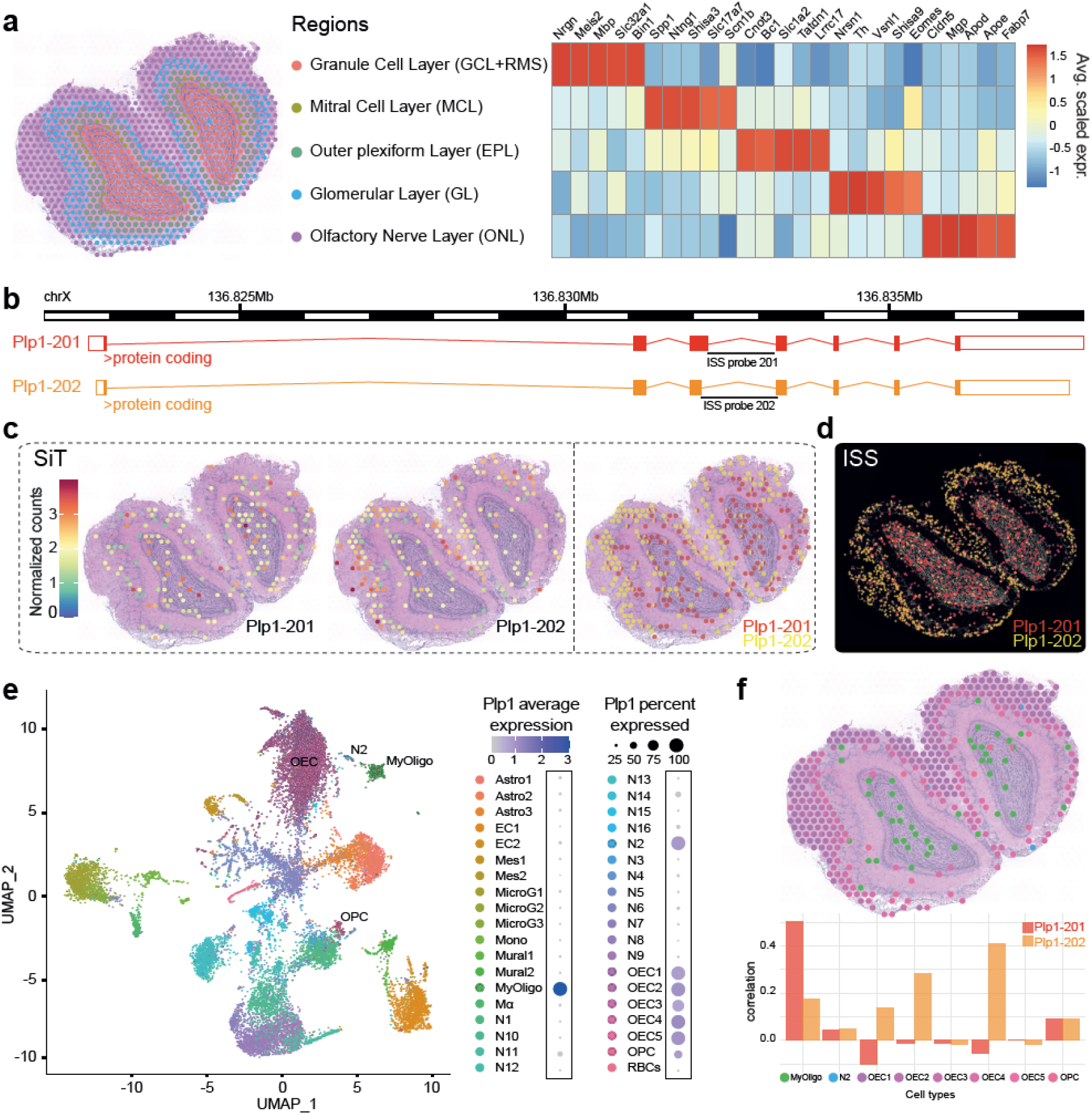
SiT reveals isoform switches in the mouse olfactory bulb. **(a)** Data-driven annotation (upper panel) of mouse olfactory bulb spatial regions through transcriptome-clustering of short-read data. Heatmap shows the expression of prominent marker genes for each region. **(b)** Exonic structure of the different Plp1 isoforms detected by SiT (mm10 build). **(c,d)** Expression of Plp1 isoforms detected by SiT **(c)** and ISS **(d). (e)** Uniform Manifold Approximation and Projection (UMAP) representation of the MOB single cell dataset from Tepe et al ^27^. The dot plot on the right indicates Plp1 expression per cell type. **(f)** Spatial spot deconvolution of cell types with high/prominent Plp1 expression (upper panel). Each dot corresponds to a pie graph indicating cell type composition in this spot. Per spatial spot correlation observed between deconvolution score and Plp1 isoform expression (lower panel). Results show that Plp1 is predominantly expressed by olfactory ensheathing cells (OEC) in the olfactory nerve layer (ONL) and by myelinating-oligodendrocytes (MyOligo) in the granule cell layer (GCL).

Myosin Light Chain 6 (*Myl6*), codes for the non-phosphorylatable alkali light chain component of the hexameric Myosin motor protein, that has been shown to be involved in neuronal migration and synaptic remodeling in immature and mature neurons^22,23^. *Myl6* produces two main polypeptides of same size, that differ just in five of the last nine carboxy terminal amino acids: the non-muscle isoform Myl6-206 (Lc17a) and the smooth-muscle isoform Myl6-201 (Lc17b)^24^. Our data revealed a high expression of Myl6-201 in the granule cell layer while Myl6-206 is preferentially expressed in the olfactory nerve and mitral cell layer (Supplementary Fig.4).

Proteolipid protein 1 (*Plp1*), a gene involved in severe pathologies associated with CNS dysmyelination^25^, demonstrated a clear regional difference in isoform expression between the inner granule cell layer, expressing full Plp1-201 (PLP) isoform and the outer regions of the olfactory nerve layer, expressing preferentially the truncated Plp1-202 (DM20) isoform (Fig.2b-c, Supplementary Fig.5). SiT allows to clearly identify 35 codons that are exclusively present in PLP and to quantify the PLP/DM20 splicing balance in a spatial context, a balance that has been shown to be implicated in Pelizaeus-Merzbacher disease^26^.

We validated the differential regional isoform expression of *Myl6* and *Plp1* using an independent hybridization-based technology, *in situ* sequencing (ISS), on a tissue section from another individual (Fig.2d, Supplementary Fig.4).

### Cell type inference reveals origin of *Plp1* isoforms

Each spatially barcoded spot typically captures transcripts from multiple cells. Single cell RNA-seq data allow to deconvolute the transcriptional signal into the likely constituent cell types of the spot, and to associate specific cell type(s) to spatial isoform expression data. We used a previously published mouse olfactory bulb single cell RNA-seq dataset^27^ to perform a deconvolution strategy based on the identification of pairwise cell correspondence^28^ (Fig.2e). This approach identified the myelinating-oligodendrocyte (MyOligo) cell type within the granule cell layer as the predominant origin of the *Plp1* standard isoform and the olfactory ensheathing cell (OEC) within the olfactory nerve layer as the predominant producer of the truncated *Plp1* isoform DM20 (Fig.2f).

### Deep sequencing of coronal brain sections

SiT was then applied to two 50 μm spaced adjacent coronal brain sections (CBS1, CBS2). We generated short- and long-read data following the same protocol as for the MOB sample^15^. The higher complexity of the spatial coronal brain section libraries motivated a deeper sequencing than for the MOB (Fig.1b). We generated a total of 174 and 287 million long reads with three and eight Oxford Nanopore flow cells, for CBS1 and CBS2 respectively (Supplementary Table 1). Clustering based on short-read gene expression data defined 12 anatomical regions (Fig.3a, Supplementary Fig.6) that broadly corresponded to regions in the Allen mouse brain reference atlas^29^ (Fig.3b). To assess the robustness of our method, we computed the transcriptome expression correlation between the two sections after image alignment and minimization of the spot-to-spot distance between sections (Supplementary Fig.7). We observed a Pearson correlation of 0.98 and 0.93 respectively for short-read and long-read gene-level profiles of corresponding pairs of spatial spots (Fig.3c).

**Figure 3.**
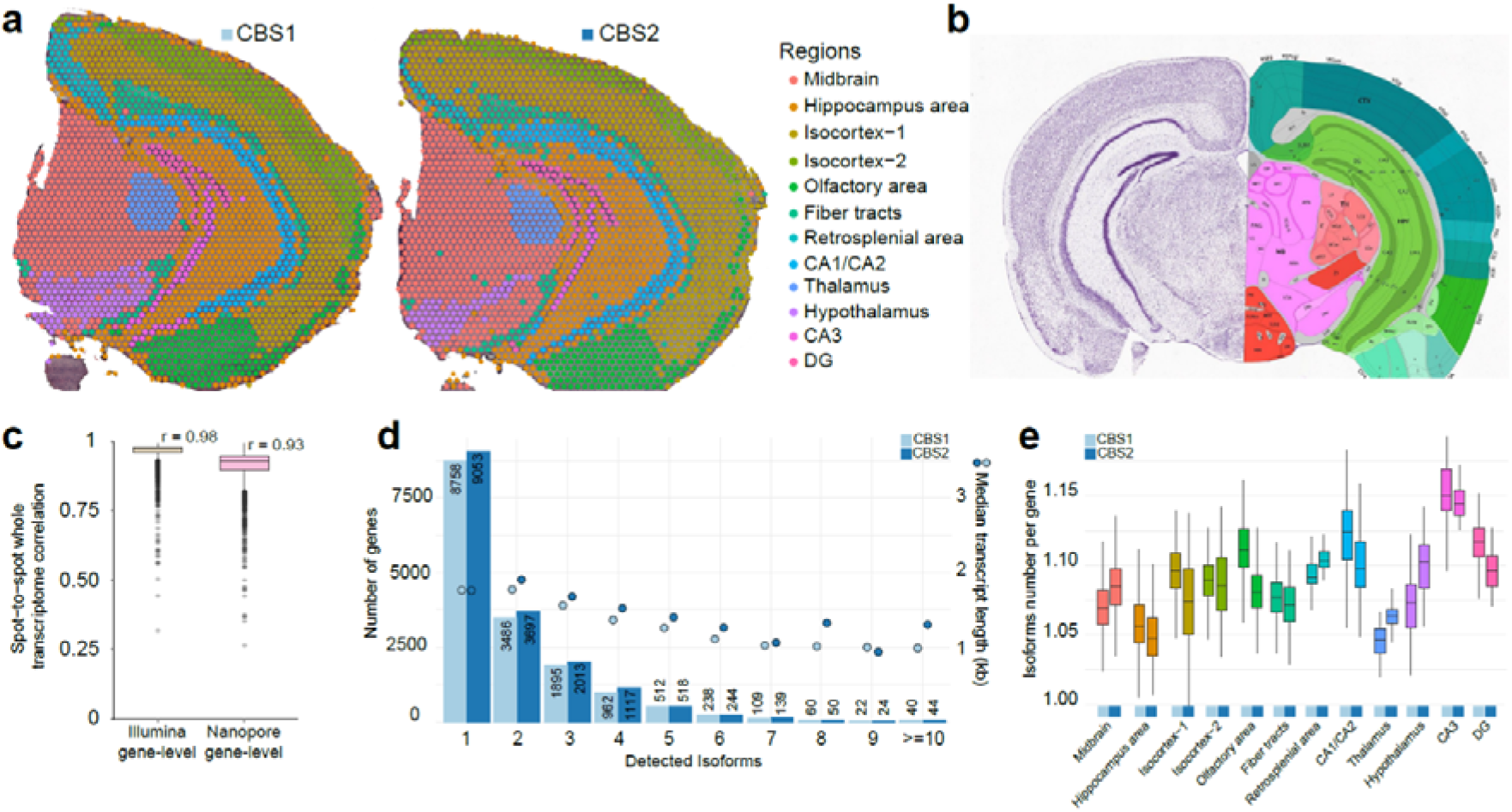
SiT robustness assessment using two coronal brain sections. **(a)** Data-driven annotation of mouse coronal brain regions through transcriptome clustering of shortread data. **(b)** Allen Mouse Brain Atlas reference map for the coronal brain section region annotation. **(c)** Gene-level short-read and long-read transcriptome correlation between corresponding spatial spots of the CBS1 and CBS2 section after image alignment and distance minimization. **(d)** Histogram showing the frequency distribution of the number of isoforms per gene for CBS1 and CBS2. The median length of transcripts is indicated for each category. **(e)** Average number of isoforms per gene detected for each spatial region.

### Regional isoform switching in coronal brain sections

We sought to identify genes with differential isoform usage across the mouse brain regions. With our stringent filters, we only retained UMIs that could be unambiguously assigned to a Gencode reference isoform (i.e., all exon-exon junctions must be observed). For CBS2, we successfully assigned 10 million molecules (UMIs) to a precise isoform, corresponding to 33,097 distinct isoforms encoded by 16,899 genes. Among those genes, we observed 9,053 (53.6%) that expressed a single isoform and 7,846 (46.4%) that expressed multiple isoforms across the tissue section (Fig.3d). We obtained a median of 3,644 UMIs for each spatially barcoded spot, corresponding to a median of 1,524 unique isoforms (Supplementary Fig.3). We noticed small variations in the number of isoforms per gene across the different brain regions and a slightly higher isoform complexity in the CA3 region of the hippocampus (Fig.3e). The numerical values for Fig.3e are also provided in Supplementary Table 3. We noticed that the number of detected genes increased with the number of spots per region. Among the multi-isoform genes, we mined for those showing a splicing pattern change across regions to decipher differences in spatial cell organization. We identified 126 and 166 significant (Bonferroni-adjusted p-value < 0.05) regional isoform switching genes in CBS1 and CBS2 respectively, out of which 61 were identified in both sections (Methods, Supplementary Table 4-5).

#### Hypothalamus expresses high level of a Snap25a isoform

For both sections, our data revealed a pronounced regional isoform switching for several genes involved in brain function (Fig.4a-b). A first example is *Snap25* which is expressed as the Snap25-202 (Snap25a) isoform in the hypothalamus, in contrast to the midbrain where Snap25-201 (Snap25b) is the predominant isoform (Fig.4c-d). Both isoforms result from inclusion/exclusion of two closely spaced sequences which encodes distinct fifth exons which results in 9 out of 206 amino acids changes between the two polypeptides, a difference that has been shown to play a role in plasticity at central synapses^30^. As previously described, Snap25a is the dominant transcript during embryonic and early postnatal development in mouse brain, while in adulthood, Snap25b becomes the dominant mRNA. Snap25a remains the dominant isoform in endocrine and neuroendocrine cells throughout life^31^. The spatial isoform expression pattern observed was confirmed by ISS (Fig.4e).

**Figure 4.**
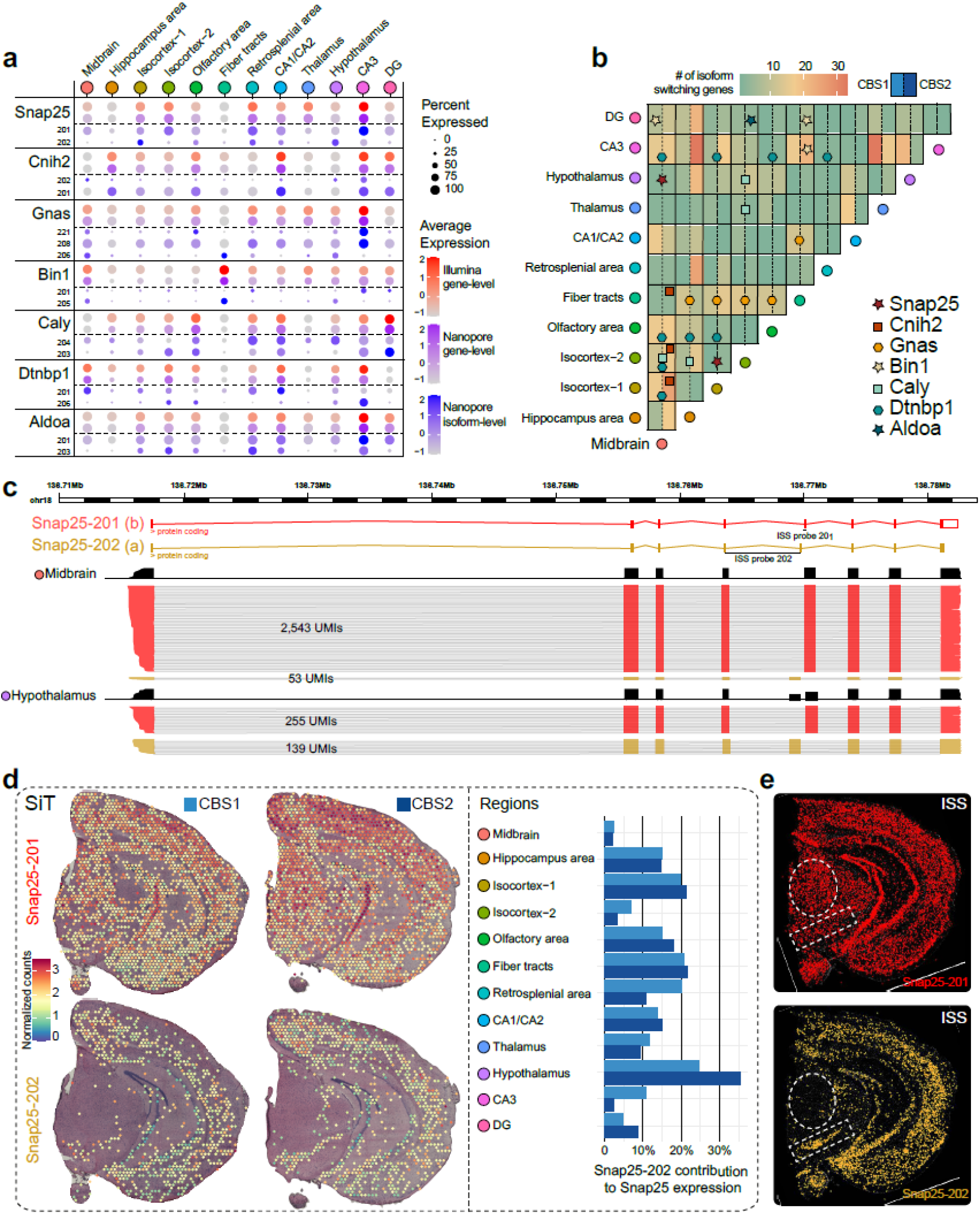
SiT reveals Snap25 isoform-switch between mouse brain regions. **(a)** Dot-plot showing the 7 most significant (adjusted p-value < 0.05) regional differences in isoform usage detected in CBS1 and CBS2 sections (CBS2 expression value is shown). **(b)** Heatmap illustrating the number of isoform-switching genes identified between 2 brain regions (same order for the horizontal and vertical axes). Genes discussed in this study are highlighted. **(c)** Snap25 isoforms (alternative exon 5) tracks and number of molecules (UMI) observed in Midbrain and Hypothalamus (CBS2). **(d)** Snap25 isoform expression in coronal brain sections revealed by SiT (left panel), Snap25-202 contribution to total Snap25 expression in brain regions in CBS1 and CBS2 (right panel). **(e)** Snap25 isoform expression validation by in situ-sequencing (ISS).

#### Dense neuronal regions and white matter express different Bin1 isoforms

A second example of regional isoform switching is *Bin1*, which belongs to the Bin-Amphiphysin-Rvs167 (BAR) domain superfamily proteins. *Bin1* is involved in the regulation of membrane curvature, particularly in clathrin-coated synaptic vesicles^32^. The *Bin1* locus has been identified as a leading modulator of genetic risk in Alzheimer’s disease^33^. Our data reveals a clear regional differential *Bin1* isoform usage. Midbrain and fiber tracts express Bin1-205 (human Bin1iso9), while the Bin1-201 isoform (human Bin1iso1) was expressed in a sparse pattern including in the isocortex, and the hippocampal formation, with enrichment in the Dentate Gyrus (DG) and the CA3 region. Spatial deconvolution using the single-cell Mouse Brain Atlas dataset^34^ revealed a high correlation between Bin1-205 isoform expression and Oligodendrocytes especially the MOL1 and MOL3 subtypes, which delineates more precisely the expression of Bin1-205 (Bin1iso9) in these two subtypes of mature oligodendrocytes^35^ (Supplementary Fig.8).

#### The Gnas locus shows complex isoform expression pattern across brain regions

A third example of regional isoform switching is *Gnas*, encoded by a complex imprinted locus^36^ as the alpha-subunit of the stimulatory G protein (Gsα), an important component of the cyclic AMP signaling pathway^37^. In both coronal brain sections, we observed a high expression of *Gnas* α-L (Gnas-208) across all regions making it the most abundant *Gnas* isoform. SiT identified multiple *Gnas* isoforms such as the low expressed splice variant *Gnas* α-S (Gnas-206) present mainly in fiber tracts as well as Gnas-221, a paternally imprinted allele specific isoform, expressed in isocortex and in the CA3 region of the hippocampus, including a restricted expression in or adjacent to the posterior amygdalar nucleus (PA) region according to the Allen Mouse Brain reference atlas (Supplementary Fig.9).

SiT identified several additional differences in the regional isoform usage for an additional set of genes including *Cnih2, Caly, Dtnpb1* and *Aldoa*, which were confirmed by ISS (Supplementary Fig.10).

### SiT reveals A-to-I RNA editing mouse brain map

We next examined whether SiT enables exploration of Single Nucleotide Variation (SNV). We investigated RNA adenosine-to-inosine (A-to-I) editing events, the principal source of transcript sequence heterogeneity in the mammalian transcriptome. RNA editing has been shown to be essential for neurotransmission and other neuronal functions^38^. While other studies examined editing events on bulk samples from mouse brain^39^, or spatially resolved by ISS for a limited number of targeted editing sites^40^, none has yet provided an exhaustive spatially-resolved RNA editing map of the mouse brain.

Robust SNV calling requires substantial sequencing depth, and we therefore focused on CBS2 for this purpose (Supplementary Table 1). We explored a total of 5,817 exonic A-to-I RNA editing sites described in the literature^39,41^. To ensure high confidence calls with Nanopore long reads, we defined an *ad hoc* UMI sequencing depth and a consensus call accuracy threshold for Nanopore editing site calls by examining the agreement between long- and short-read editing site data for the same UMI (RNA molecule) for 70,225 UMIs where at least one known editing site was observed in both Illumina and Nanopore molecules (total 88,175 editing site observations). Based on the analysis shown in Supplementary Fig. 11, we retained UMIs backed by at least 3 reads with a consensus quality at the editing site greater than 6 for further analysis. This resulted in a > 99% agreement between Illumina and Nanopore editing site calls. Out of the 377,304 Nanopore editing site observations, 249,759 (66.2%) passed those filters and were used for downstream analysis (Fig.5a, Methods). Globally, we observed an A-to-I RNA editing ratio of 10.56% for 2,730 distinct editing sites covered by at least one UMI (46.9% of the 5,817 known editing sites explored, Supplementary Table 6). Interestingly, editing ratios displayed a non-uniform spatial distribution (Fig.5b). Consistent with a previous report^40^, we observed a significantly higher editing ratio in Thalamus (mean 17.6%; 749/4,247 UMIs edited) than in Fiber tracts (mean 5.4%; 657/12,063 UMIs edited). We also noticed a positive correlation between the expression levels of the A-to-I editing enzymes (adenosine deaminases, ADARs) and the editing ratios for the brain regions (Fig.5c). The same variation between distinct brain areas was independently noticed in both coronal brain sections (Supplementary Fig.12), which displays similar profiles. We observed a Pearson correlation score of 0.91 between CBS1 and CBS2 editing ratio for the 483 editing sites showing at least 20 UMIs in CBS1 and CBS2 long reads profiles (Supplementary Fig.13).

**Figure 5.**
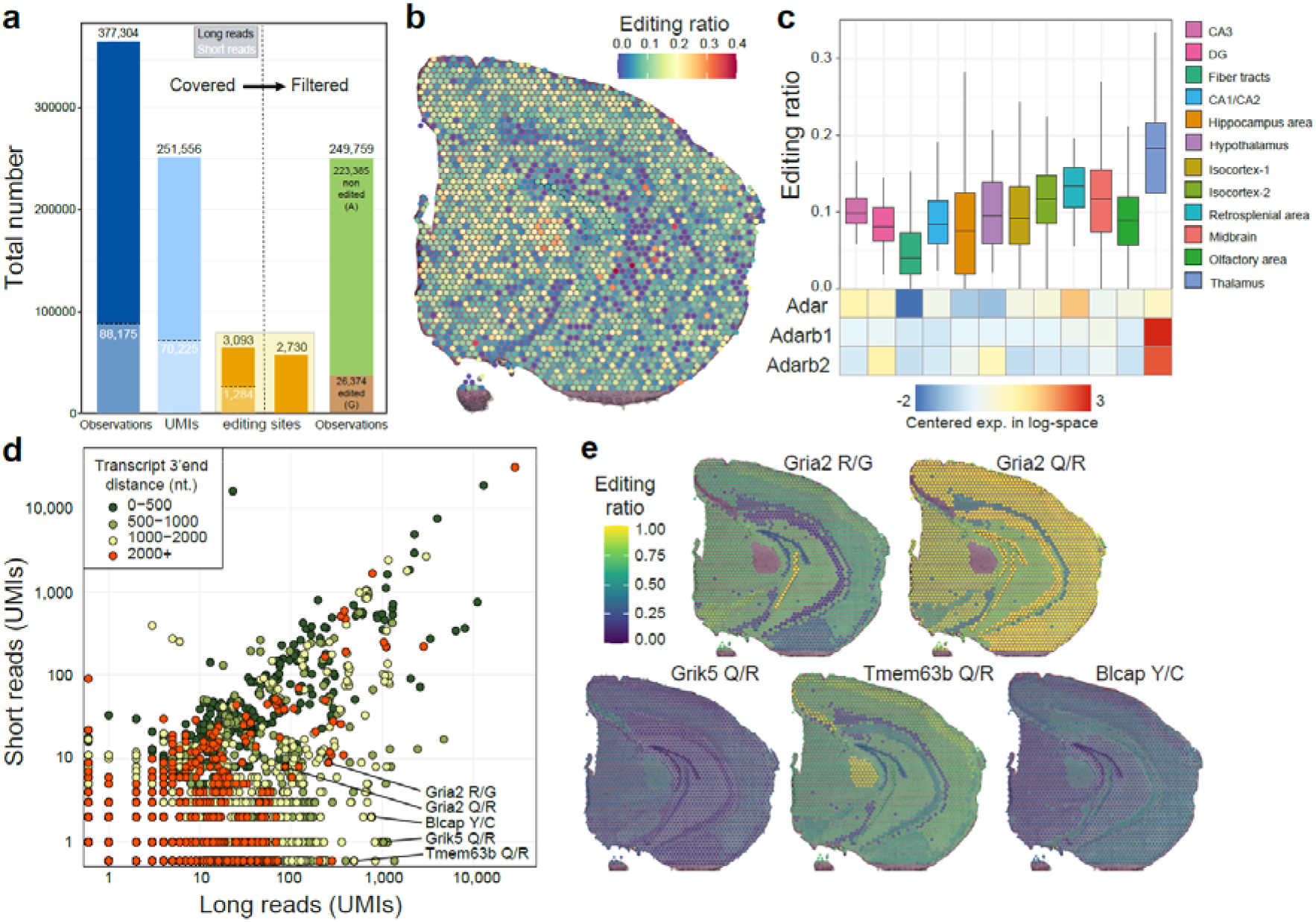
SiT reveals spatial variations in A-to-I RNA editing in the mouse brain. **(a)** Histogram showing the sum of known editing site observations on all UMIs (Observations, blue bars), the total number of UMIs covering at least one editing site (UMIs, light blue bars), the number of distinct known exonic editing sites described in the literature^39,41^ (orange bars) from long read (black numbers) and short read (white numbers) data covering editing sites before (left part) and after (right part) quality filtration of Nanopore UMIs (consensus sequence quality >=6 and UMI sequencing depth >= 3, see Supplementary Fig.11). **(b)** Spatial map of A-I editing ratios. **(c)** Editing ratio and gene-level expression of ADAR enzymes per region. **(d)** Editing site short- and long-read coverage colored by distance to transcripts 3’-end. **(e)** Spatial map of the A-I editing ratio for a selection of editing sites. Editing ratio computed as the average editing ratio of molecules observed per region.

We next sought to compare the potential of short and long read sequencing to identify RNA editing events. As expected, we observed that long reads yield more information than short reads (Fig.5d) since the long-read sequencing allowed to investigate sequence heterogeneity beyond the 3’-end editing sites. This resulted in the identification of editing variation across brain regions in several key genes of neuronal function such as *Gria2*^42^, *Grik5*^43^, *Tmem63b*^44^ or *Blcap*^45^ (Fig.5e).

AMPA receptors (AMPARs) mediate most of the fast excitatory neurotransmission in the brain and are formed from four different subunits. The transcript for the Gria2 (GluA2) subunit is known to be edited at two positions: the R/G site (mm10, chr3:80692286), which is in the ligand-binding domain where editing causes faster desensitization and recovery from desensitization, and the Q/R site (mm10, chr3:80706912) located within the channel pore, which when edited renders AMPARs virtually Ca2+-impermeable and thereby affects a key aspect of neurotransmission^46^. Consistent with previous observations^47^, we observed less editing at the R/G site (mean=55.5%, 142/256 UMIs edited) than at the Q/R site (mean=94.2%, 81/86 UMIs edited). We also observed regional differences in R/G site editing with low editing levels in subregions of the hippocampus (DG granule cell layer: mean=36.4%, 4/11 edited UMIs; CA1/CA2: mean=20.5%, 8/39 UMIs edited) and high editing levels in the isocortex region (mean=67.5%, 79/117 UMIs edited) (Fig.5e).

## DISCUSSION

Here we present SiT, the first transcriptome-wide approach to explore isoform expression and sequence heterogeneity in a tissue context. We demonstrate the ability of SiT to identify differential isoform usage across brain regions. We further validated the ability of SiT to detect A-to-I RNA editing events in full-length transcript sequences. In the mouse olfactory bulb, we demonstrated a clear isoform switch of *Plp1*, a gene coding for the major myelin protein in the nervous system, between the outer regions of the olfactory nerve layer and the inner granule cell layer. In coronal brain sections we showed differential isoform usage for several other key genes, including *Snap25, Bin1* or *Gnas*. We further established the spatial pattern of A-to-I RNA editing events in an unbiased manner, to provide for the first time a map of global editing ratios in the adult mouse brain and show that SiT enables investigation of individual editing sites in a spatial context. We opted for Nanopore sequencing since this technology has a far lower cost per read than PacBio sequencing. However SiT can easily be adapted for PacBio sequencing.

One limitation of SiT is that only 26 - 40% of the Nanopore reads both matched the reference genome and were assigned to a valid spatial barcode and a UMI for the three sections while 63 - 90% of the Illumina reads met those criteria. The principal reasons for the high fraction of unassigned Nanopore reads are^15^: (i) elimination of low quality Nanopore reads where no spatial barcode and UMI can be identified. Further improvements in Nanopore sequencing accuracy will increase assignment efficiency significantly, (ii) About 20% of the sequences do not contain the RT primer with the cell barcode and the UMI and are discarded. Optimization of the library preparation should significantly increase assignment efficiency, for instance by further depletion of unwanted cDNA sequences that do not contain a polyA tail. (iii) While our Illumina-guided cell barcode and UMI assignment strategy is highly accurate^15^, the efficiency of UMI assignment to Nanopore reads depends on the sequencing saturation of the short read dataset which was 71%, 73% and 93% for the CBS1, CBS2 and MOB section respectively. Nanopore reads with UMIs that were not found in the Illumina dataset cannot be assigned and are lost. Development of an Illumina-free assignment strategy will help recovering molecules missed during Illumina sequencing and increase the efficiency of UMI assignment. Another limitation of SiT is that the resolution is so far limited to the resolution provided by the Visium technology. Several alternatives were recently described that increase the resolution of spatial transcriptomics (Slide-SeqV2^48^, DBiT-seq^49^). SiT can be readily adapted to those approaches which generate full length cDNA. A last point, also shared with all single-cell approaches, regards the limitations on the number and integrity of captured mRNA molecules. Only half of the captured molecules contain all exons of a Gencode isoform and are unambiguously attributed to a full-length isoform. The rest corresponds to truncated cDNA, likely derived from degraded RNA (Supplementary Fig.1). Further optimizations of the Visium workflow aiming at reduced RNA degradation prior to cDNA synthesis and at more efficient capture of mRNA molecules should address those limitations.

Our results show that a throughput of 100 million long reads, now obtained routinely with one PromethION flow cell, is sufficient to explore the spatial landscape of transcript isoform expression in a typical Visium™ experiment. Increased sequence accuracy and throughput of the nanopore technology will further benefit to the SiT approach. We anticipate that such progresses should make SiT more and more applicable in many different environments, including in different clinical contexts. A straightforward application would certainly be in cancerology to resolve spatially the expression of pathological isoforms (e.g., fusion transcripts) and cancer mutations in order to better characterize the heterogeneity of tumour biopsies. SiT could thus contribute to the advent of more efficient therapeutic avenues.

SiT expands the spatial transcriptomics toolbox to long-read based exploration of transcript isoforms and SNVs, such as RNA editing or somatic mutations. These observations have thus far escaped detection due to the limitations of conventional short-read sequencing approaches. In combination with whole-brain molecular maps^50^, we show here how this approach offers a new opportunity to understand the spatial and molecular organization of complex organs.

The SiT methodology is based on commercially available reagents and enables a deepened investigation of the isoform landscape, including studies of imprinting, fusion transcripts, and SNV expression in a spatial context. SiT will enable a better description of complex transcriptomes. As such, it provides an important additional resource to enrich existing Cell Atlas initiatives, as illustrated here through the on-line resource of mouse brain that we provide (https://www.isomics.eu/).

## METHODS

### Mouse Brain Samples

Olfactory bulbs were isolated from C57BL/6 mice (>2 months old), snap-frozen in Isopentane (Sigma-Aldrich) and embedded in cold optimal cutting temperature (OCT, Sakura) before sectioning. The left hemisphere was isolated from an C57BL/6J (8-12 weeks old) mouse and processed in the same way. Olfactory bulbs from two different individuals were used for the Visium and ISS experiments, whereas the same sample of the left hemisphere was used for both methods.

### 10x Genomics Visium experiments

The Visium Spatial Tissue Optimization Slide & Reagent kit (10xGenomics, Pleasanton, CA, USA) was used to optimize permeabilization conditions for mouse brain tissue. Two coronal sections of the left hemisphere (IDs: CBS1 and CBS2) and one section of olfactory bulb (ID: “MOB”) were processed according to the manufacturer’s protocol. Spatially barcoded full-length cDNA was generated using Visium Spatial Gene Expression Slide & Reagent kit (10xGenomics) following the manufacturer’s protocol. Tissue permeabilization was performed for 6 and 9 min (CBS1, CBS2) and 12 min (MOB). cDNA amplification was conducted with 12 (CBS) and 17 (MOB) cycles. A fraction of each cDNA library was used for Nanopore sequencing, whereas 10 μl was used in the 10xGenomics Visium library preparation protocol of fragmentation, adapter ligation, and indexing. The libraries were sequenced on a NextSeq500 (Illumina), with 28 bases from read 1 and 91 from read 2, and at a depth of 253, 217, and 210 million reads for MOB, CBS1, and CBS2 samples, respectively. The raw sequencing data was processed with a pre-launch of the Space Ranger pipeline (10xGenomics) and mapped to the mm10 genome assembly.

### Oxford Nanopore sequencing

Nanopore sequencing of libraries prepared with cDNA from the 10xGenomics workflow yields 20-50% reads without the 3’ adapter sequence and thus lacks the spatial barcode and UMI. To deplete such DNA, we initially selected for cDNA that contains a biotinylated 3’ primer. 10 ng of the 10xGenomics Visium PCR product were amplified for 5 cycles with 5’-AAGCAGTGGTATCAACGCAGAGTACAT-3’ and 5’ Biotine-AAAAACTACACGACGCTCTTCCGATCT 3’. Excess biotinylated primers were removed by 0.55x SPRIselect (Beckman Coulter) purification and the biotinylated cDNA (in 40 μl EB, Qiagen) was bound to 15 μl 1x SSPE washed Dynabeads™ M-270 Streptavidin beads (Thermo) in 10 μl 5x SSPE for 15 min at room temperature on a shaker. Beads were washed twice with 100 μl 1x SSPE and once with 100 μl EB. The beads were suspended in 100 μl 1x PCR mix and amplified for 8 cycles with the primers NNNAAGCAGTGGTATCAACGCAGAGTACAT and NNNCTACACGACGCTCTTCCGATCT to generate enough material (1 – 2 μg) for Nanopore sequencing library preparation. To deplete small fragments which are typically of little interest for transcript isoform analysis (cDNA from degraded RNA, ribosomal RNAs), small cDNA (< 1 kB) was depleted with a 0.5x SPRI select purification. If fragments between 0.5 and 1 kB need to be retained, SPRIselect concentration should be increased to 0.8x. Nanopore sequencing libraries were prepared with the LSK-109 or LSK-110 kit from Oxford nanopore (1 μg cDNA) following the instructions from the manufacturer. PromethION flow cells were loaded with 200 ng libraries each. PCR amplifications for Nanopore library preparations were made with Kapa Hifi Hotstart polymerase (Roche Sequencing Solutions): initial denaturation, 3 min at 95°C; cycles: 98°C for 30 sec, 64°C for 30 sec, 72°C for 5 min; final elongation: 72°C for 10 min, primer concentration was 1 μM.

### Oxford Nanopore data processing

Nanopore reads were processed according to the scNaUmi-seq protocol^15^ with slight modifications. Briefly, to eliminate reads that originate from chimeric cDNA generated during library preparation, we initially scanned reads for internal (> 200 nt from end) Template Switching Oligonucleotide (TSO, AAGCAGTGGTATCAACGCAGAGTACAT) and 3’ adapter sequences (CTACACGACGCTCTTCCGATCT) flanked by a poly(T) (poly(T)-adapter). When two adjacent poly(T)-adapters, two TSOs or one TSO in proximity of a poly(T)-adapter were found, the read was split into two separate reads. Next the reads were scanned for poly(A/T) tails and the 3’ adapter sequence to define the orientation of the read and strand-specificity. Scanned reads were then aligned to *Mus musculus* mm10 with minimap2 v2.17 in spliced alignment mode. Spatial BCs and UMIs were then assigned to nanopore reads using the strategy and software previously described for single cell libraries^15^. SAM records for each spatial spot and gene were grouped by UMI after removal of low-quality mapping reads (mapqv=0) and potentially chimeric reads (terminal Soft/Hard-clipping of > 150 nt). A consensus sequence per molecule (UMI) was computed depending on the number of available reads for the UMI using the *ComputeConsensus* sicelore-2.0 method. For molecules supported by more than two reads (RN > 2), a consensus sequence was computed with SPOA^51^ using the sequence between the end of the TSO (SAM Tag: TE) and the base preceding the polyA sequence (SAM Tag: PE). Quality values for consensus nucleotides were assigned as −10*log_10_(n Reads not conform with consensus nucleotide / n Reads total), maximum set to 20. Consensus cDNA sequences were aligned to the *Mus musculus* mm10 build with minimap2 v2.17 in spliced alignment mode. SAM records matching known genes were analyzed for matching Gencode vM24 transcript isoforms (same exon makeup). To assign a UMI to a Gencode transcript, we required a full match between the UMI and the Gencode transcript exon-exon junction layout authorizing a two-base margin of added or lacking sequences at exon boundaries, to allow for indels at exon junctions and imprecise mapping by minimap2. Detailed statistics of each step of Nanopore read processing are provided in Supplementary Table 1.

### Data analysis and count matrices storage

Raw gene expression matrices generated by Space Ranger were processed using R/Bioconductor (version 4.0.2) and the Seurat R package (version 3.9.9). We created Seurat objects for each of the three samples (MOB, CBS1 and CB2) with different assays for the analysis as follows: (i) “Spatial” containing gene-level raw short read data from the Space Ranger output, (ii) “ISOG” containing the gene-level Nanopore long read data, (iii) “ISO” containing isoform-level transcript information where only the molecules where all exons are observed are kept, (iv) “JUNC” containing each individual exon-exon junction observation per isoform, and (v) “Atol” containing exonic editing sites from the RADAR database (mm9 UCSC liftover to mm10) and from the Licht et al., 2019, study, for which we observed at least one UMI in our dataset. The “Atol assay stored non edited UMI count (@counts slot), edited UMI count (@data slot), and the editing ratio (@scale.data slot) per editing site.

### 10xGenomics Visium data-driven annotation of anatomical regions

The Spatial assay was normalized with *SCTransform* using standard parameters. The first 30 principal components of the assay were used for UMAP representation and clustering (resolution = 0.4). Brain regions defined by clustering were assigned to known anatomical regions based on the Allen Mouse Brain Atlas. Spot clustering was similar between short and long read data (Supplementary Fig.14). As short read data contains more UMIs per spot, our different gene markers representations are based on short read data.

### Spatial spot deconvolution

For MOB, spatial spots were deconvoluted using SPOTlight (version 0.1.4) and signature genes identified from main Tepe et al., 2018 ^27^ WT samples *Plp1* expresser cell types (mean normalized expression > 1) identified using Seurat *FindAllMarkers* (logfc.threshold = 0.25, min.pct = 0.1). Cell types contributing to at least 8% were selected and SPOTlight deconvolution scores were used for correlation computation with *Plp1* isoforms expression. The same approach was performed for coronal brain section (CBS1 and CBS2) using Zeisel et al., 2018 ^34^ external dataset (mean expression > 1 UMI/cell).

### Differential splicing detection

Seurat *FindMarkers* function (logfc.threshold = 0.25, test.use = “wilcox”, min.pct = 0.1) was used to detect genes showing at least 2 isoforms as markers of different brain regions using the Nanopore isoform-level “ISO” assay. Results were filtered for non-majority isoforms, i.e., not the isoform showing the highest bulk expression, requiring Bonferonni-adjusted p.value ≤ 0.05.

### Coronal brain sections transcriptome correlation

Images were masked then Aligned using *MaskImages* and *AlignImages* STUtility^52^ R package (version 1.0.0) function before we computed and minimized the physical distance between spots to defined pair of spots showing the smallest distance between sections. We then computed the whole transcriptome correlation per pair of spots using *cor.test* function from Stats R package using genelevel short-read (Spatial assay) and long-read (ISOG assay) UMI count matrices.

### Long-read calibration for high-confidence RNA editing call

To only keep high confidence base calls, the Nanopore data were filtered by exploring the percentage of agreement between both sequencing methods as a function of long read number (RN) per molecule and Nanopore consensus base quality value (Supplementary Table 7). Long-read molecules having a minimum read number of 3 (MINRN=3) and a base quality value at the editing position of 6 (MINQV=6) were chosen to be of sufficient quality for editing sites calling using the *SNPMatrix* sicelore-2.0 method.

### Editing ratios

Samples CBS2 and MOB were used for calculating global editing ratios. To test the significance of our findings, resampling of capture-spots across the sample were performed. Observed editing ratios per spot were kept and each spot was randomly assigned a region-label from the pool of original labels without replacement 10k times. A normal distribution was fitted to the simulated editing ratios to calculate the probability of observing a value equal to, or more extreme, than the observed value. Region-level individual editing site editing ratio was computed as the total edited molecules divided by the total molecules observed at the editing site position within each region.

### *In situ* sequencing validation

Ten μm cryosections of the olfactory bulb and coronal sections of the left hemisphere were placed on SuperFrost Plus microscope slides (ThermoFisher Scientific), stored at −80 °C and shipped on dry ice to CARTANA for library preparation, probe hybridization, probe ligation, rolling circle amplification, and fluorescence labeling using the HS Library Preparation Kit (P/N 1110) and for the *in-situ* sequencing using the ISS kit (P/N 3110) and sequential imaging using a 20x objective. The result table of the spatial coordinates of each molecule of all targets together with the reference DAPI image per sample were provided by CARTANA.

## Supporting information

Supplementary Figure 1. Size distribution of cDNA

Supplementary Figure 2. Comparison between Illumina and nanopore gene-level data

Supplementary Figure 3. Isoform UMI counts and number of isoforms per capture spot

Supplementary Figure 4. Mouse Olfactory Bulb (MOB) Myl6 isoform expression

Supplementary Figure 5. Plp1 coverage plot in the 5 spatial regions defined in mouse olfactory bulb (MOB)

Supplementary Figure 6. Spatial annotation of CBS2 brain regions driven by short-read data

Supplementary Figure 8. Bin1 isoform expression in coronal brain sections

Supplementary Figure 9. Gnas isoform expression in coronal brain sections

Supplementary Figure 10. Caly, Cnih2, Dtnbp1, and Aldoa isoform expression in CBSs

Supplementary Figure 11. Percentage of agreement between long-read and short-read data for editing sites that are detected by both approaches

Supplementary Figure 12. Resampling of global editing ratios and test statistics

Supplementary Figure 13. Correlation of editing ratios between coronal brain sections CBS1 and CBS2 (a) and between short and long-reads (b)

Supplementary Figure 14. Coronal Brain Section (CBS2) spatial spots classification after SCTransform normalization

## Data availability

All relevant data have been deposited in Gene Expression Omnibus under accession number GSE153859 (https://www.ncbi.nlm.nih.gov/geo/query/acc.cgi?acc=GSE153859).

## Code availability

All custom software used is available on Github (https://github.com/ucagenomix/sicelore). R figures and analysis scripts are available on Github (https://github.com/ucagenomix/SiT). Seurat object .rds files for the three samples are available on demand and interactively browsable in the dedicated SiTx R Shiny application accessible at https://www.isomics.eu.

## Acknowledgements

This project was supported by Institut National contre le Cancer (PLBIO2018-156), FRM (DEQ20180339158), the Inserm Cross-cutting Scientific Program HuDeCA 2018, the National Infrastructure France Génomique (Commissariat aux Grands Investissements, ANR-10-INBS-09-03, ANR-10-INBS-09-02), the 3IA@cote d’azur (<u>ANR-19-P3IA-0002</u>), the Swedish Research Council, Swedish Foundation for Strategic Research, European Union’s H2020 Research and Innovation Program under grant agreement no. 874656 (discovAIR), Conseil départemental 06, Knut and Alice Wallenberg Foundation (2018.0172), Erling-Persson Family Foundation (HDCA), and Science for Life Laboratory. We would like to thank the National Genomics Infrastructure (NGI) Sweden for providing infrastructure support. We thank Ludvig Bergenstråhle and Alma Andersson for advice and helpful discussions.

## Contributions

K.T., A.M., R.W. performed the experiments. K.L, J.B. analyzed the data. P.B., R.W. and J.L. supervised the research. All authors contributed to the writing of the manuscript.

## References

1. Baralle, F. E. & Giudice, J. Alternative splicing as a regulator of development and tissue identity. Nature Reviews Molecular Cell Biology vol. 18 437–451 (2017).

2. Su, C. H., Dhananjaya, D. & Tarn, W. Y. Alternative splicing in neurogenesis and brain development. Frontiers in Molecular Biosciences vol. 5 12 (2018).

3. Herbrechter, R., Hube, N., Buchholz, R. & Reiner, A. Splicing and editing of ionotropic glutamate receptors: a comprehensive analysis based on human RNA-Seq data. Cell. Mol. Life Sci. 2021 7814 78, 5605–5630 (2021).

4. Lipscombe, D. & Lopez Soto, E. J. Alternative splicing of neuronal genes: new mechanisms and new therapies. Current Opinion in Neurobiology vol. 57 26–31 (2019).

5. Yang, Y., Okada, S. & Sakurai, M. Adenosine-to-inosine RNA editing in neurological development and disease. https://doi.org/10.1080/15476286.2020.1867797 18, 999–1013 (2021).

6. Sapiro, A. L. et al. Illuminating spatial A-to-I RNA editing signatures within the Drosophila brain. Proc. Natl. Acad. Sci. 116, 2318–2327 (2019).

7. Costa Cruz, P. H. & Kawahara, Y. Rna editing in neurological and neurodegenerative disorders. in Methods in Molecular Biology vol. 2181 309–330 (Humana Press Inc., 2021).

8. Asp, M., Bergenstråhle, J. & Lundeberg, J. Spatially Resolved Transcriptomes—Next Generation Tools for Tissue Exploration. BioEssays 1900221 (2020) doi:10.1002/bies.201900221.

9. Rodriques, S. G. et al. Slide-seq: A scalable technology for measuring genome-wide expression at high spatial resolution. Science (80-.). 363, 1463–1467 (2019).

10. Stickels, R. R. et al. Sensitive spatial genome wide expression profiling at cellular resolution. doi:10.1101/2020.03.12.989806.

11. Liu, Y. et al. High-Spatial-Resolution Multi-Omics Atlas Sequencing of Mouse Embryos via Deterministic Barcoding in Tissue. SSRN Electron. J. (2019) doi:10.2139/ssrn.3466428.

12. Chen, A. et al. Title: Large field of view-spatially resolved transcriptomics at nanoscale resolution Short title: DNA nanoball stereo-sequencing. doi:10.1101/2021.01.17.427004.

13. Hagemann-Jensen, M. et al. Single-cell RNA counting at allele and isoform resolution using Smart-seq3. Nat. Biotechnol. 38, 708–714 (2020).

14. Gupta, I. et al. Single-cell isoform RNA sequencing characterizes isoforms in thousands of cerebellar cells. Nat. Biotechnol. 36, 1197–1202 (2018).

15. Lebrigand, K., Magnone, V., Barbry, P. & Waldmann, R. High throughput error corrected Nanopore single cell transcriptome sequencing. Nat. Commun. 11, 1–8 (2020).

16. Volden, R. et al. Improving nanopore read accuracy with the R2C2 method enables the sequencing of highly multiplexed full-length single-cell cDNA. Proc. Natl. Acad. Sci. U. S. A. 115, 9726–9731 (2018).

17. Sakamoto, Y., Sereewattanawoot, S. & Suzuki, A. A new era of long-read sequencing for cancer genomics. J. Hum. Genet. 65, 3 (2020).

18. Amarasinghe, S. L. et al. Opportunities and challenges in long-read sequencing data analysis. Genome Biol. 2020 211 21, 1–16 (2020).

19. Mincarelli, L., Uzun, V., Rushworth, S. A., Haerty, W. & Macaulay, I. C. Combined single-cell gene and isoform expression analysis in haematopoietic stem and progenitor cells. doi:10.1101/2020.04.06.027474.

20. Joglekar, A. et al. A spatially resolved brain region- and cell type-specific isoform atlas of the postnatal mouse brain. Nat. Commun. 12, 1–16 (2021).

21. Ståhl, P. L. et al. Visualization and analysis of gene expression in tissue sections by spatial transcriptomics. Science vol. 353 78–82 (2016).

22. Kneussel, M. & Wagner, W. Myosin motors at neuronal synapses: drivers of membrane transport and actin dynamics. Nat. Rev. Neurosci. 14, 233–247 (2013).

23. Vallee, R. B., Seale, G. E. & Tsai, J.-W. Emerging roles for myosin II and cytoplasmic dynein in migrating neurons and growth cones. Trends Cell Biol. 19, 347 (2009).

24. Chen, P. et al. The expression and functional activities of smooth muscle myosin and non-muscle myosin isoforms in rat prostate. J. Cell. Mol. Med. 22, 576–588 (2018).

25. Nave, K. A. Myelination and support of axonal integrity by glia. Nature vol. 468 244–252 (2010).

26. Regis, S., Grossi, S., Corsolini, F., Biancheri, R. & Filocamo, M. PLP1 gene duplication causes overexpression and alteration of the PLP/DM20 splicing balance in fibroblasts from Pelizaeus-Merzbacher disease patients. Biochim. Biophys. Acta - Mol. Basis Dis. 1792, 548–554 (2009).

27. Tepe, B. et al. Single-Cell RNA-Seq of Mouse Olfactory Bulb Reveals Cellular Heterogeneity and Activity-Dependent Molecular Census of Adult-Born Neurons. Cell Rep. 25, 2689–2703.e3 (2018).

28. Elosua-Bayes, M., Nieto, P., Mereu, E., Gut, I. & Heyn, H. SPOTlight: seeded NMF regression to deconvolute spatial transcriptomics spots with single-cell transcriptomes. Nucleic Acids Res. 49, e50–e50 (2021).

29. Sunkin, S. M. et al. Allen Brain Atlas: An integrated spatio-temporal portal for exploring the central nervous system. Nucleic Acids Res. 41, D996 (2013).

30. Irfan, M. et al. SNAP-25 isoforms differentially regulate synaptic transmission and long-term synaptic plasticity at central synapses. Sci. Reports 2019 91 9, 1–14 (2019).

31. Bark, C. I., Hahn, K. M., Ryabinin, A. E. & Wilson, M. C. Differential expression of SNAP-25 protein isoforms during divergent vesicle fusion events of neural development. Proc. Natl. Acad. Sci. U. S. A. 92, 1510–1514 (1995).

32. Zhang, B. & Zelhof, A. C. Amphiphysins: Raising the BAR for synaptic vesicle recycling and membrane dynamics. Traffic vol. 3 452–460 (2002).

33. Chapuis, J. et al. Increased expression of BIN1 mediates Alzheimer genetic risk by modulating tau pathology. Mol. Psychiatry 18, 1225–1234 (2013).

34. Zeisel, A. et al. Molecular Architecture of the Mouse Nervous System. Cell 174, 999–1014.e22 (2018).

35. De Rossi, P. et al. Predominant expression of Alzheimer’s disease-associated BIN1 in mature oligodendrocytes and localization to white matter tracts. Mol. Neurodegener. 11, (2016).

36. Bastepe, M. The GNAS Locus: Quintessential Complex Gene Encoding Gsα, XLαs, and other Imprinted Transcripts. Curr. Genomics 8, 398–414 (2008).

37. Turan, S. & Bastepe, M. GNAS Spectrum of Disorders. Current Osteoporosis Reports vol. 13 146–158 (2015).

38. Behm, M. & Öhman, M. RNA Editing: A Contributor to Neuronal Dynamics in the Mammalian Brain. Trends in Genetics vol. 32 165–175 (2016).

39. Licht, K. et al. A high resolution A-to-I editing map in the mouse identifies editing events controlled by pre-mRNA splicing. Genome Res. 29, 1453–1463 (2019).

40. Lundin, E. et al. Spatiotemporal mapping of RNA editing in the developing mouse brain using in situ sequencing reveals regional and cell-type-specific regulation. BMC Biol. 18, 1–15 (2020).

41. Ramaswami, G. & Li, J. B. RADAR: A rigorously annotated database of A-to-I RNA editing. Nucleic Acids Res. 42, D109 (2014).

42. Salpietro, V. et al. AMPA receptor GluA2 subunit defects are a cause of neurodevelopmental disorders. Nat. Commun. 2019 101 10, 1–16 (2019).

43. Chew, L. J. et al. Characterization of the Rat GRIK5 Kainate Receptor Subunit Gene Promoter and Its Intragenic Regions Involved in Neural Cell Specificity. J. Biol. Chem. 276, 42162–42171 (2001).

44. Wu, D. et al. Distant coupling between RNA editing and alternative splicing of the osmosensitive cation channel Tmem63b. J. Biol. Chem. 295, 18199 (2020).

45. Schulz, R. et al. Transcript- and tissue-specific imprinting of a tumour suppressor gene. Hum. Mol. Genet. 18, 118 (2009).

46. Pachernegg, S., Münster, Y., Muth-Köhne, E., Fuhrmann, G. & Hollmann, M. GluA2 is rapidly edited at the Q/R site during neural differentiation in vitro. Front. Cell. Neurosci. 9, 69 (2015).

47. Wen, W., Lin, C.-Y. & Niu, L. R/G editing in GluA2Rflop modulates the functional difference between GluA1 flip and flop variants in GluA1/2R heteromeric channels. Sci. Reports 2017 71 7, 1–15 (2017).

48. Stickels, R. R. et al. Highly sensitive spatial transcriptomics at near-cellular resolution with Slide-seqV2. Nat. Biotechnol. 39, 313–319 (2021).

49. Y, L. et al. High-Spatial-Resolution Multi-Omics Sequencing via Deterministic Barcoding in Tissue. Cell 183, 1665–1681.e18 (2020).

50. Ortiz, C. et al. Molecular atlas of the adult mouse brain. Sci. Adv. 6, (2020).

51. Vaser, R., Sović, I., Nagarajan, N. & Šikić, M. Fast and accurate de novo genome assembly from long uncorrected reads. Genome Res. 27, 737–746 (2017).

52. Bergenstråhle, J., Larsson, L. & Lundeberg, J. Seamless integration of image and molecular analysis for spatial transcriptomics workflows. BMC Genomics 21, (2020).

