## Supplementary Figure 1. Size distribution of cDNA for "The spatial landscape of gene expression isoforms in tissue sections"

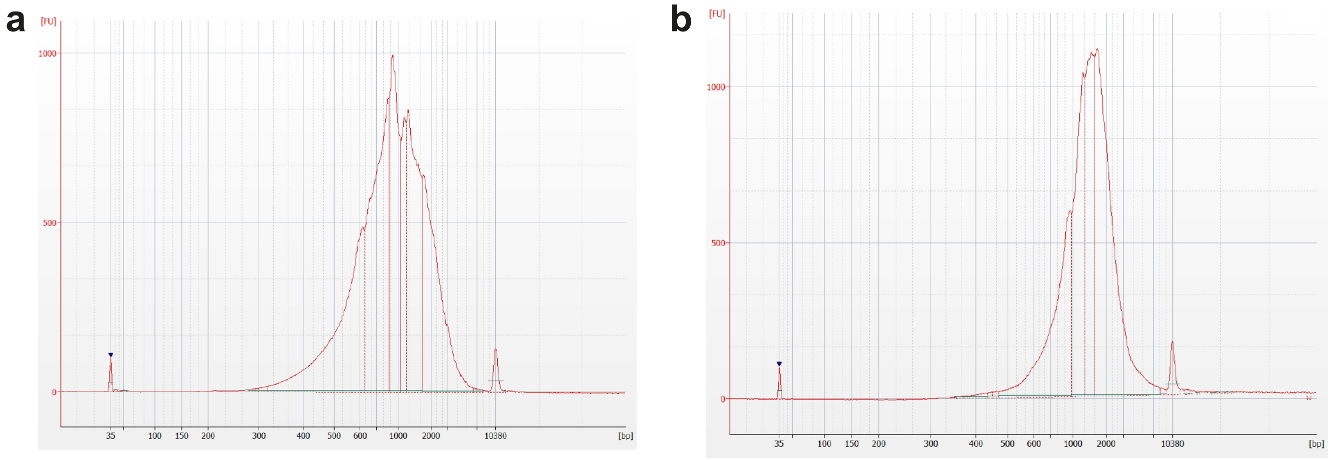


**Supplementary Figure 1. Size distribution of cDNA.** Bioanalyer profiles showing the size distribution of cDNA (CBS1 sample) used for Illumina library preparation (**a**) and for Nanopore sequencing (**b**). For Nanopore sequencing very small cDNA (< 1 kb) was partially depleted (see methods section for details).
