## Supplementary Figure 2. Comparison between Illumina and nanopore gene-level data for "The spatial landscape of gene expression isoforms in tissue sections"

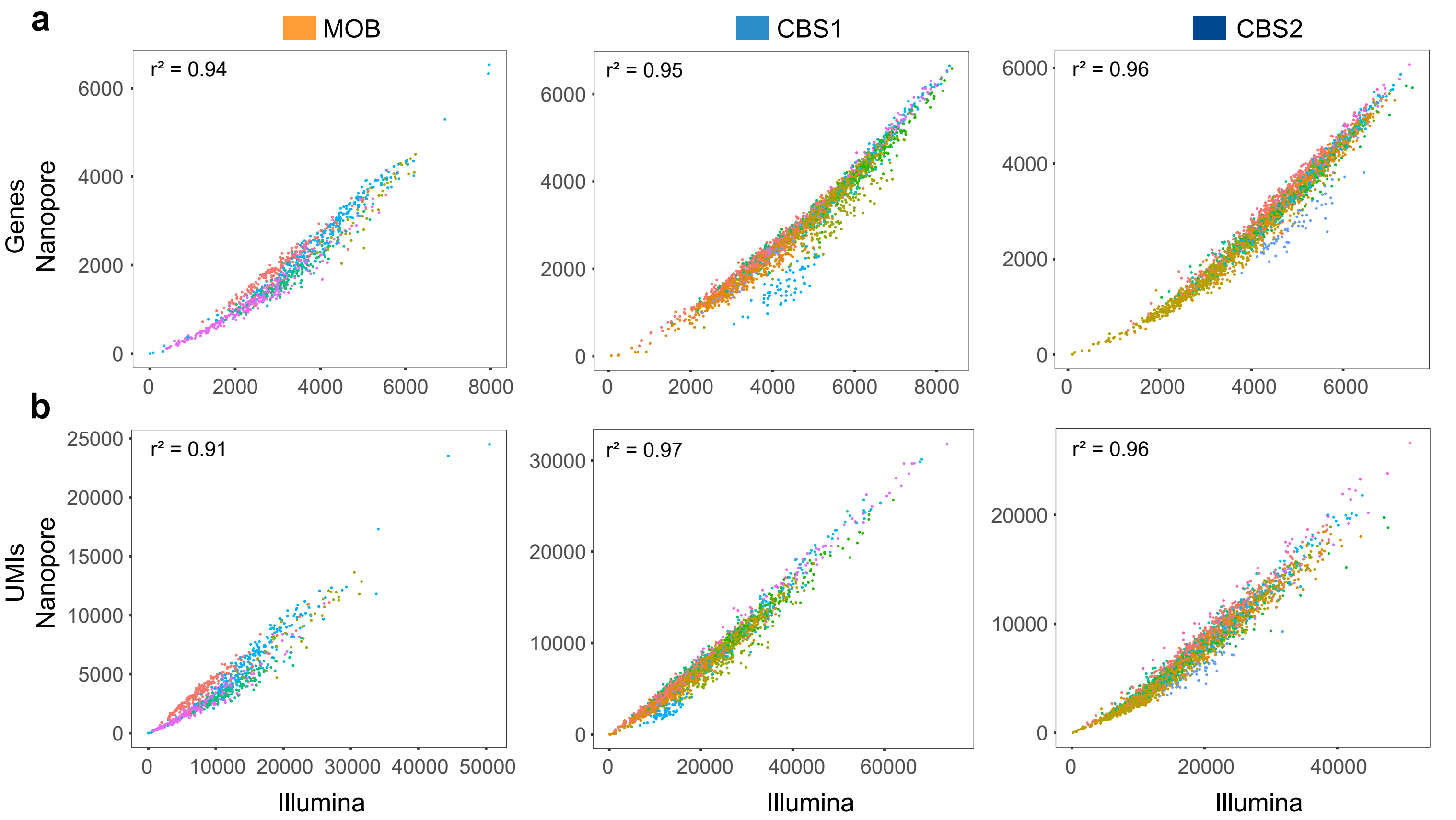


**Supplementary Figure 2. Comparison between Illumina and nanopore gene-level data.** Correlation of (**a**) Number of genes detected per capture-spot and (**b**) number of UMIs per capture-spot identified with Illumina and Nanopore sequencing. Dots are colored according to the different spatial regions defined in Fig.2a for MOB, and Fig.3a for CBS1 and CBS2.
