## Supplementary Figure 3. Isoform UMI counts and number of isoforms per capture spot for "The spatial landscape of gene expression isoforms in tissue sections"

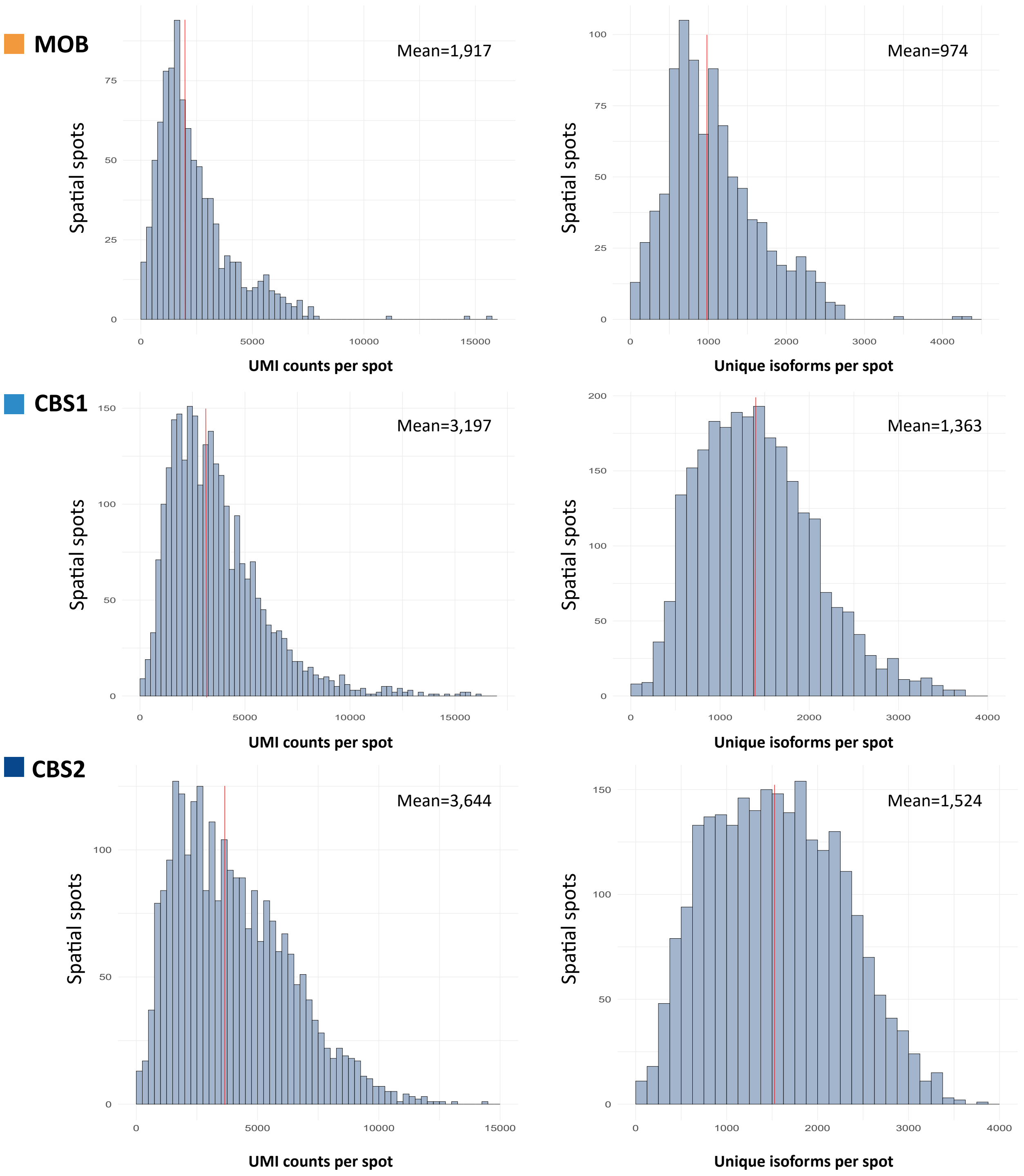


**Supplementary Figure 3. Isoform UMI counts and number of isoforms per capture spot.** Left panels show the number of UMIs assigned to a transcript isoform per capture spot. Right panels show the number of distinct transcript isoforms detected per spot. Means are indicated by red vertical lines.
