## Supplementary Figure 4. Mouse Olfactory Bulb (MOB) Myl6 isoform expression for "The spatial landscape of gene expression isoforms in tissue sections"

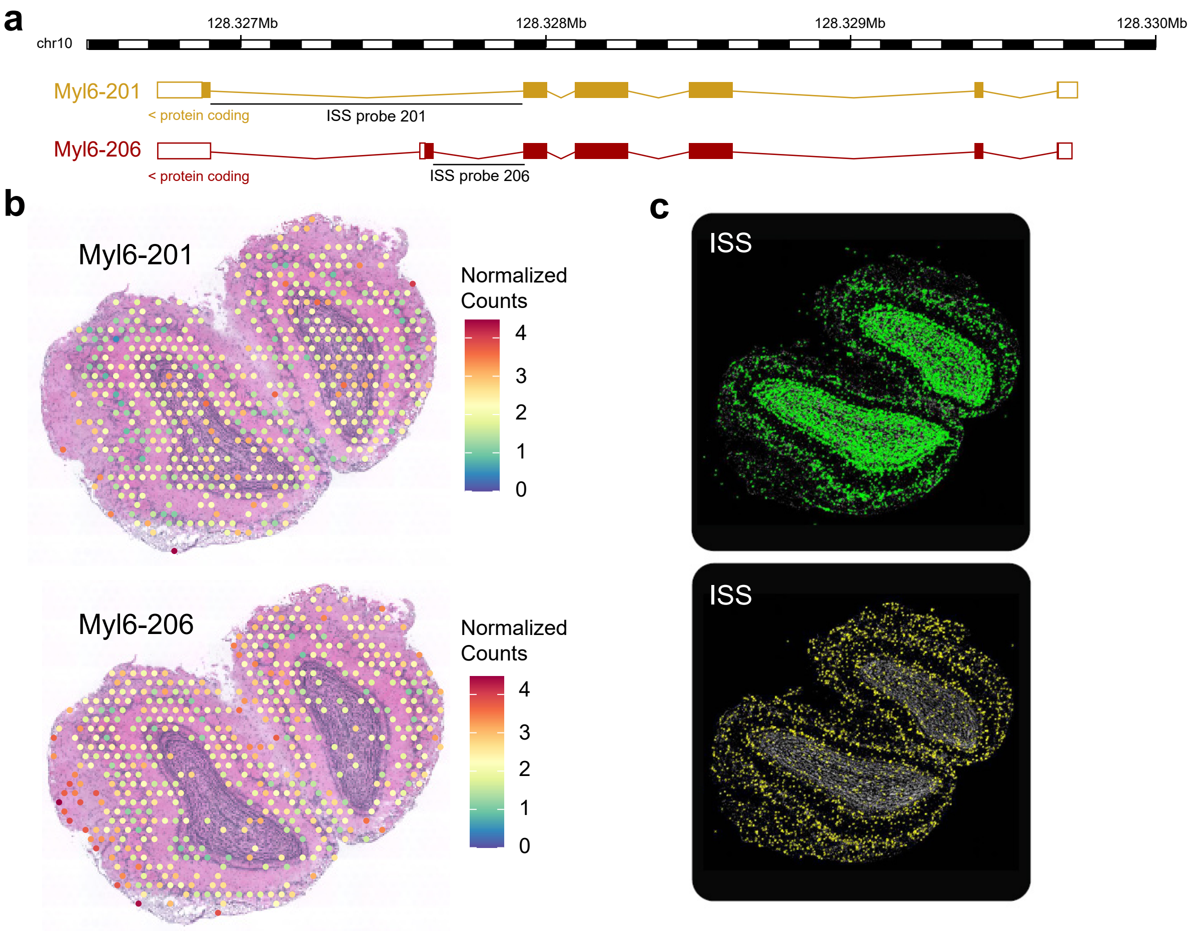


**Supplementary Figure 4. Mouse Olfactory Bulb (MOB) Myl6 isoform expression.** (**a**) Schematic view of *Myl6* gene locus (mm10 build coordinates). (**b,c**) Expression of *Myl6* isoforms detected by SiT (b) and ISS (c). ISS isoform specific probes are indicated in panel (a).
