## Supplementary Figure 5. Plp1 coverage plot in the 5 spatial regions defined in mouse olfactory bulb (MOB) for "The spatial landscape of gene expression isoforms in tissue sections"

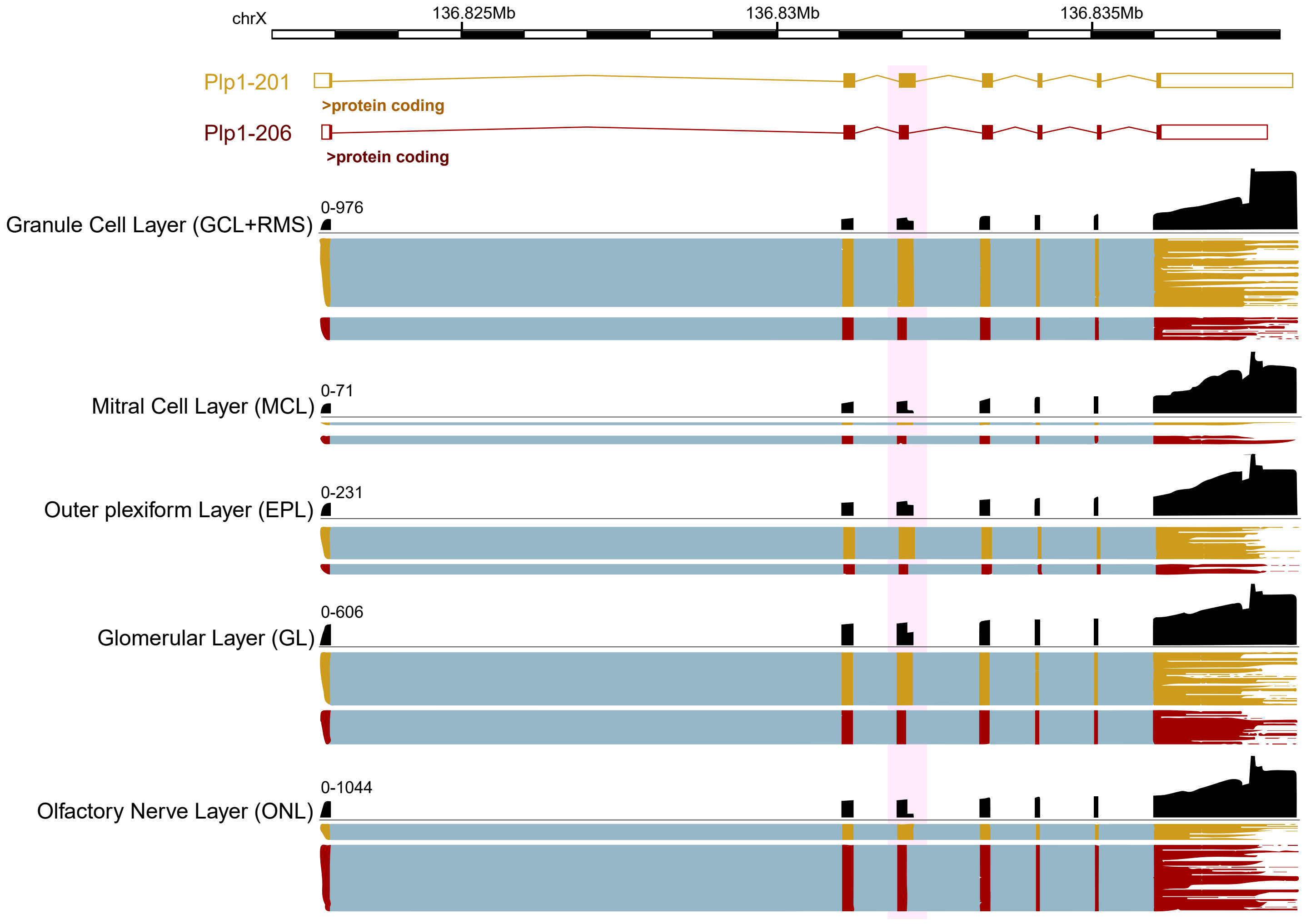


**Supplementary Figure 5. Plp1 coverage plot in the 5 spatial regions defined in mouse olfactory bulb (MOB).**
