## Supplementary Figure 6. Spatial annotation of CBS2 brain regions driven by short-read data for "The spatial landscape of gene expression isoforms in tissue sections"

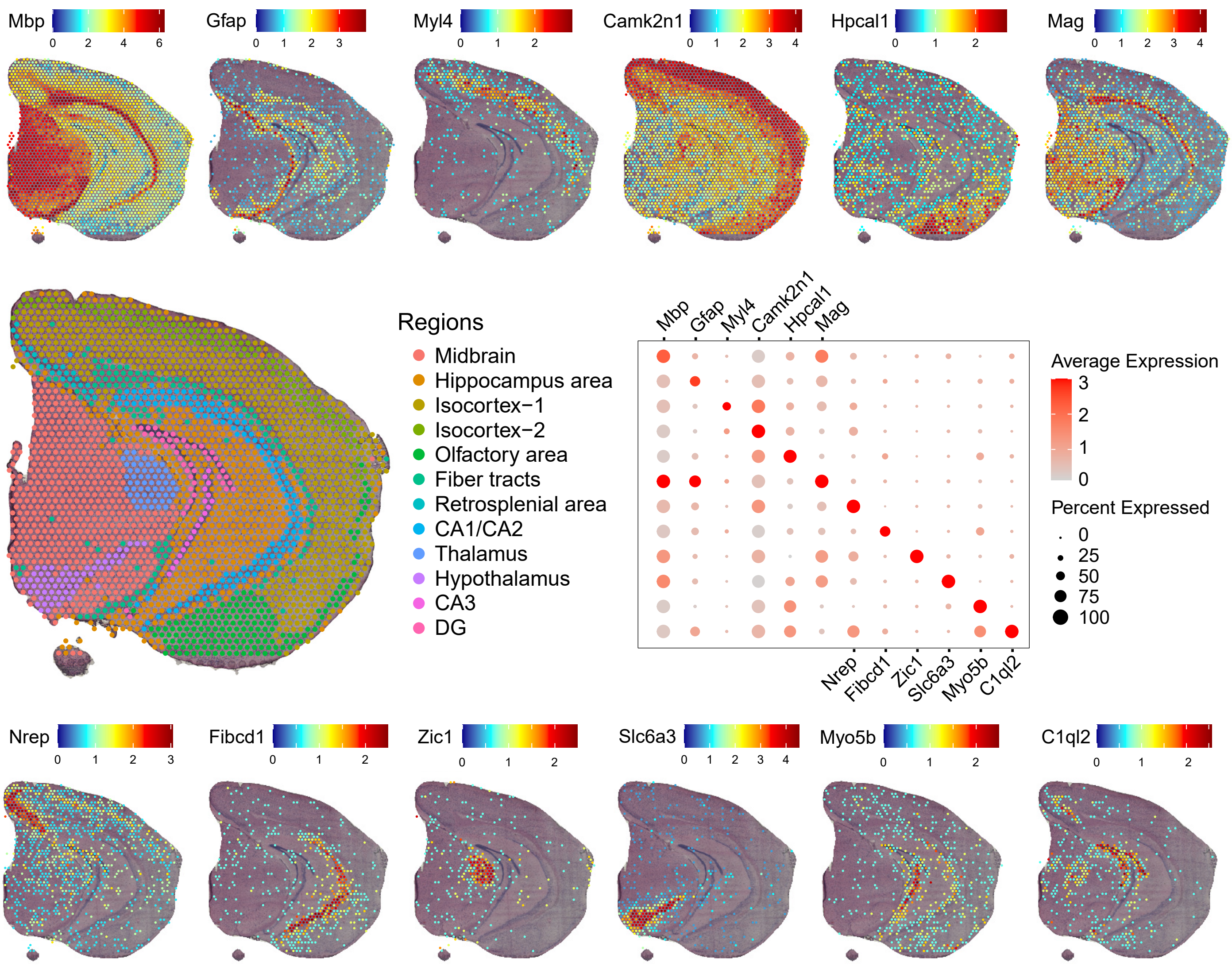


**Supplementary Figure 6. Spatial annotation of CBS2 brain regions driven by short-read data.** Spatial maps (upper and lower panels) and bubble plots (middle right panel) show the expression of the most prominent marker genes for each of the 12 annotated brain regions.
