## Supplementary Figure 7 for "The spatial landscape of gene expression isoforms in tissue sections"

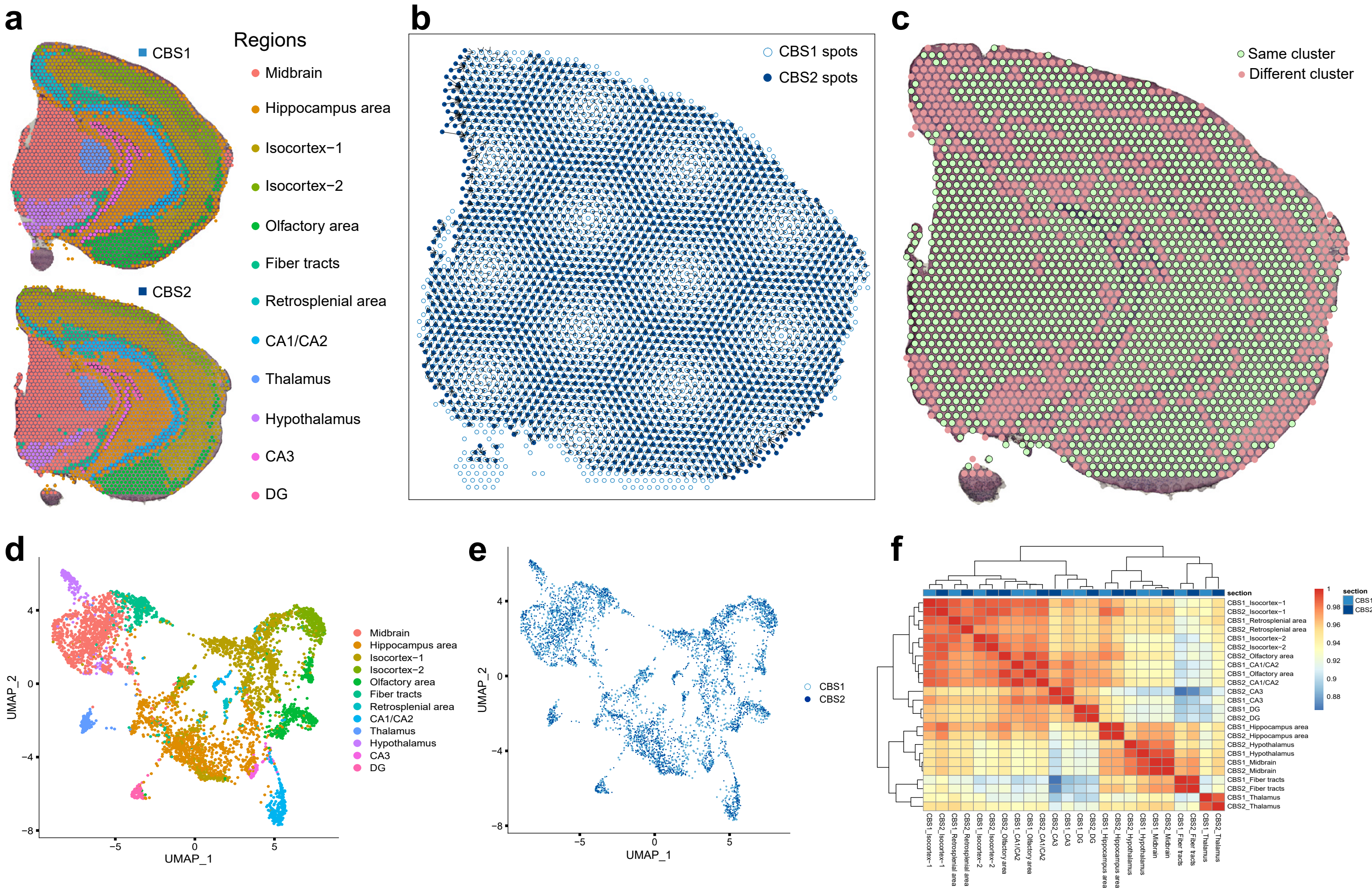


**Supplementary Figure 7.** (**a**) Data-driven annotation of the 2 coronal brain sections through transcriptome clustering of short-read data. (**b**) Correspondance between pairs of spatial spots from CBS1 and CBS2 after image alignment and distance minimization. (**c**) Visualization of pairs of spots belonging to same and different annotated regions in the two coronal brain sections. UMAP plot representation of CBS1 and CBS2 after multi-section Seurat data integration using Nanopore isoform-level assay (ISO) colored by cluster labels (**d**) and sections of origin (**e**). (**f**) Heatmap of Pearson correlation coefficient (r) between cells grouped by sections and by regions.
