## Supplementary Figure 8. Bin1 isoform expression in coronal brain sections for "The spatial landscape of gene expression isoforms in tissue sections"

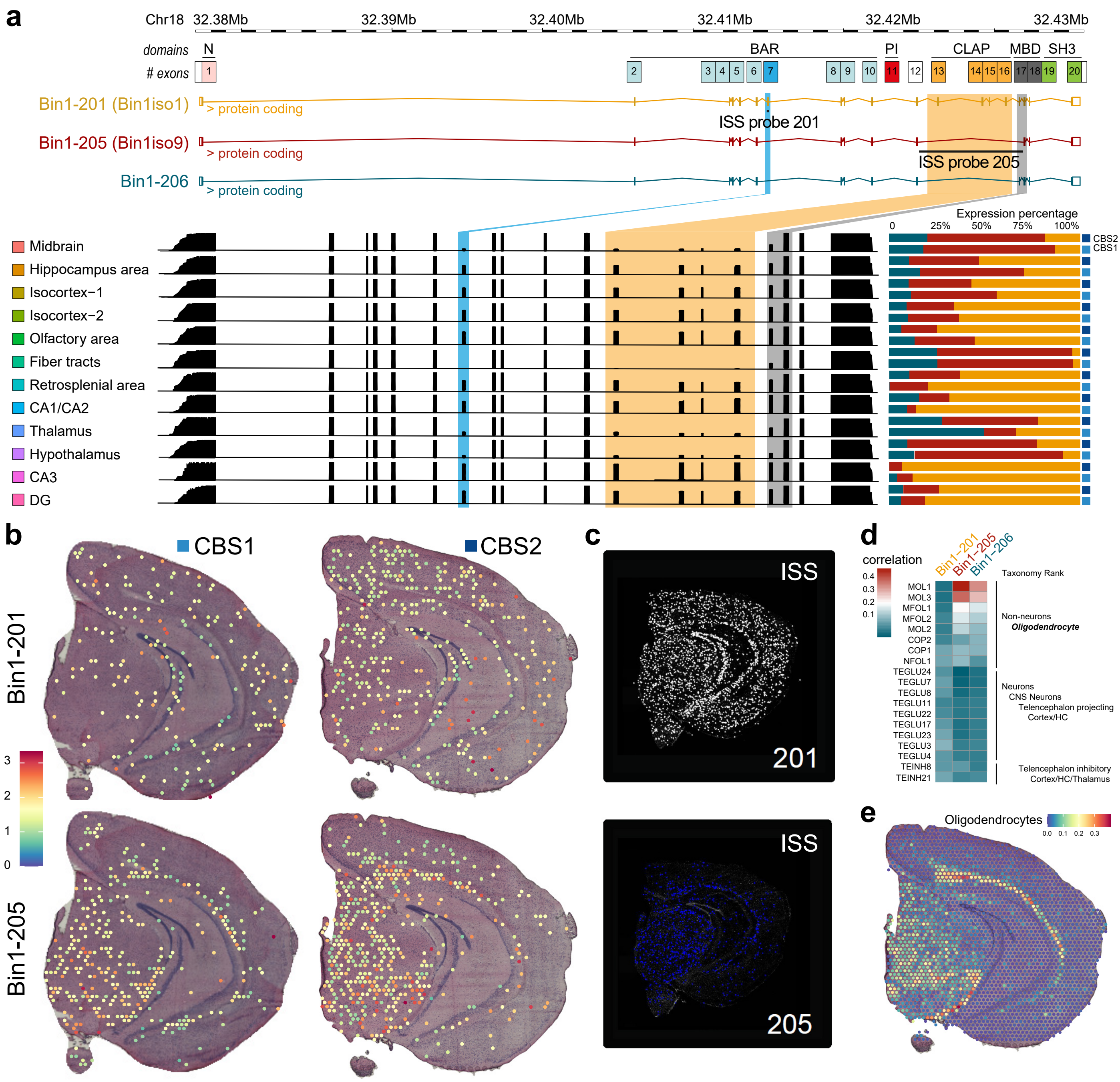


**Supplementary Figure 8**. **Bin1 isoform expression in coronal brain sections.** (**a**) Exonic structure of the different *Bin1* isoforms that are detected in CBS (upper panel); *Bin1* coverage plot in the 12 distinct CBS regions (lower left panel) and contribution of each isoform to total *Bin1* expression (lower right panel). Detection of Bin1-201 and Bin1-205 expression using SiT (**b**) and in situ sequencing (ISS) (**c**). Bin1-201 stands for human Bin1iso1 that includes exons 7 and exons 13-14-15-16-17 (CLAP domain). Bin1-205 stands for human Bin1iso9 that excludes exons 7 and CLAP domain. ISS was performed using a tissue section from another individual. (**d**) Pearson correlation coefficient (r) between *Bin1* isoform expression and the SpotLight deconvolution score for the prominent cell types that express *Bin1*. Data were derived from Zeisel et al. (2018). (**e**) SpotLight deconvolution score for Oligodendrocytes taxonomy rank across CBS2 section.
