## Supplementary Figure 9. Gnas isoform expression in coronal brain sections for "The spatial landscape of gene expression isoforms in tissue sections"

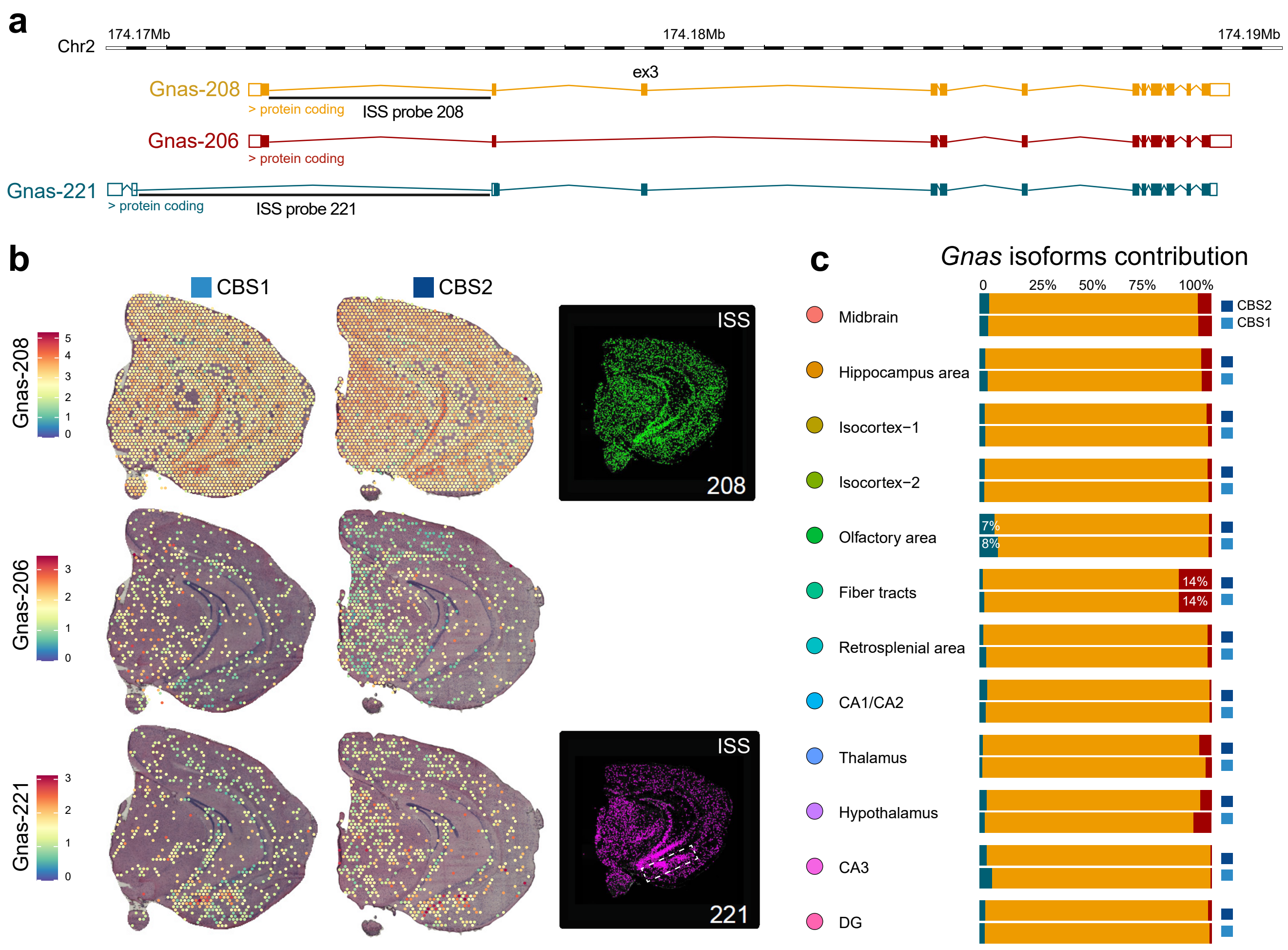


**Supplementary Figure 9. *Gnas* isoform expression in coronal brain sections.** (**a**) Exonic structure of the different ***Gnas*** isoforms that are detected in CBS. Detection of Gnas-206, Gnas-208 and Gnas-221 by SiT (**b**) and by ***in-situ*** sequencing (ISS). ISS was performed using a tissue section from another individual. (**c**) Contribution of each ***Gnas*** isoform to total Gnas expression.
