## Supplementary Figure 10. Caly, Cnih2, Dtnbp1, and Aldoa isoform expression in CBSs for "The spatial landscape of gene expression isoforms in tissue sections"

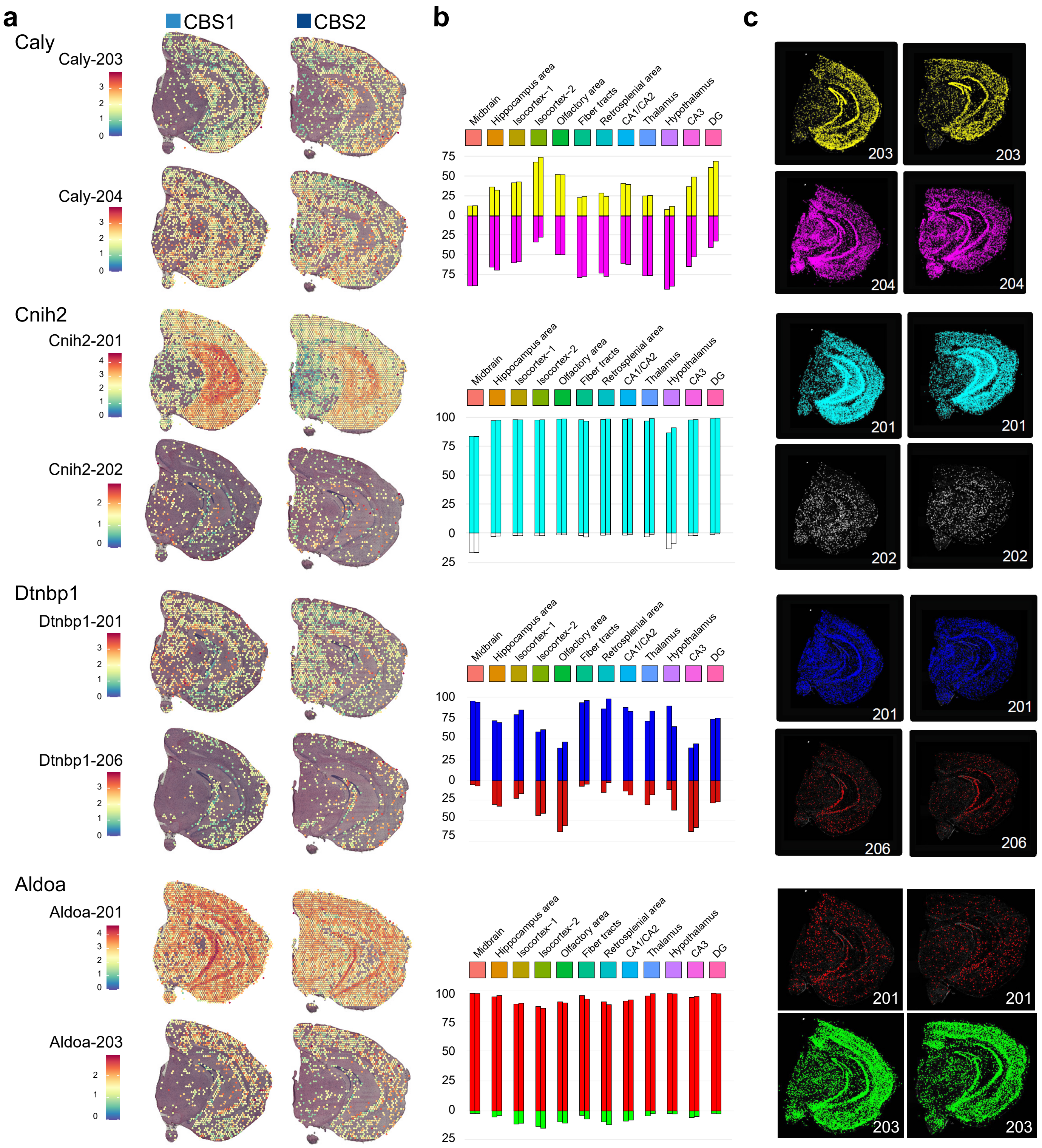


**Supplementary Figure 10. *Caly*, *Cnih2*, *Dtnbp1*, and *Aldoa* isoform expression in CBS.** (**a**) Spatial maps of isoforms expression detected by SiT. Expression data was normalized on the isoform level to highlight differences in expression across regions. Only capture spots with non-zero counts are displayed. (**b**) Quantification of associated isoforms by brain region for CBS1(left bars) and CBS2 (right bars). (**c**) *In-situ* sequenceing (ISS) with probes that target the individual transcript isoforms.
