## Supplementary Figure 11. Percentage of agreement between long-read and short-read data for editing sites that are detected by both approaches for "The spatial landscape of gene expression isoforms in tissue sections"

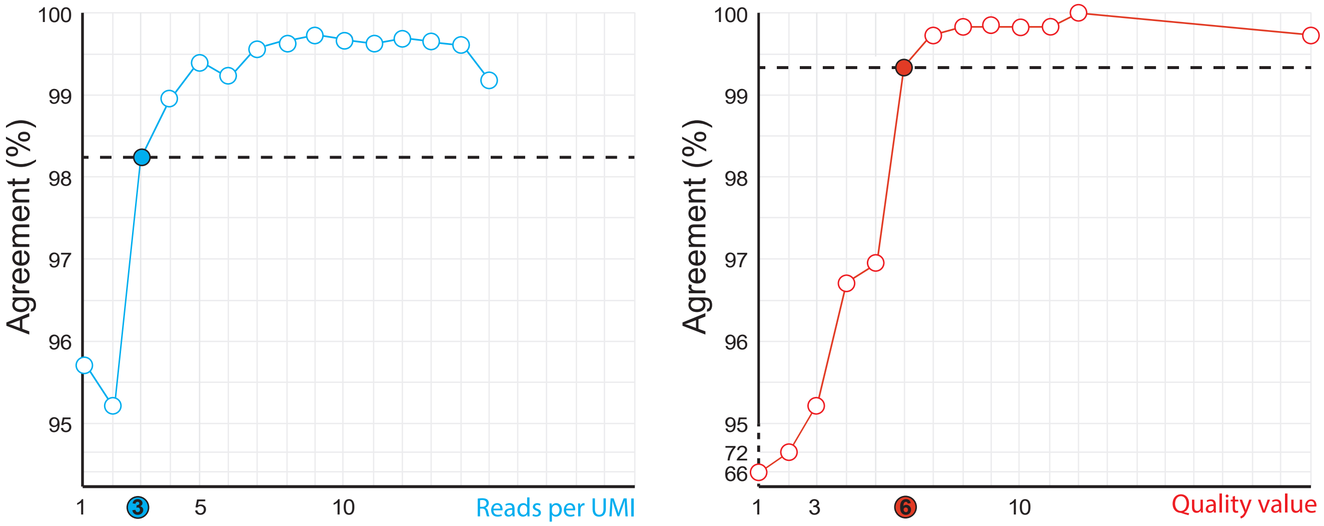


**Supplementary Figure 11. Percentage of agreement between long-read and short-read data for editing sites that are detected by both approaches.** The percentage of agreement between the two sequencing approaches for the same molecule (UMI) is plotted as a function of Nanopore read numbers per UMI (left plot) and Nanopore consensus base quality value (right plot). Quality values for consensus nucleotides were assigned as -10*log10(n Reads not conform with consensus nucleotide / n Reads total). The highlighted thresholds were used for editing site calling with Nanopore reads.
