## Supplementary Figure 12. Resampling of global editing ratios and test statistics for "The spatial landscape of gene expression isoforms in tissue sections"

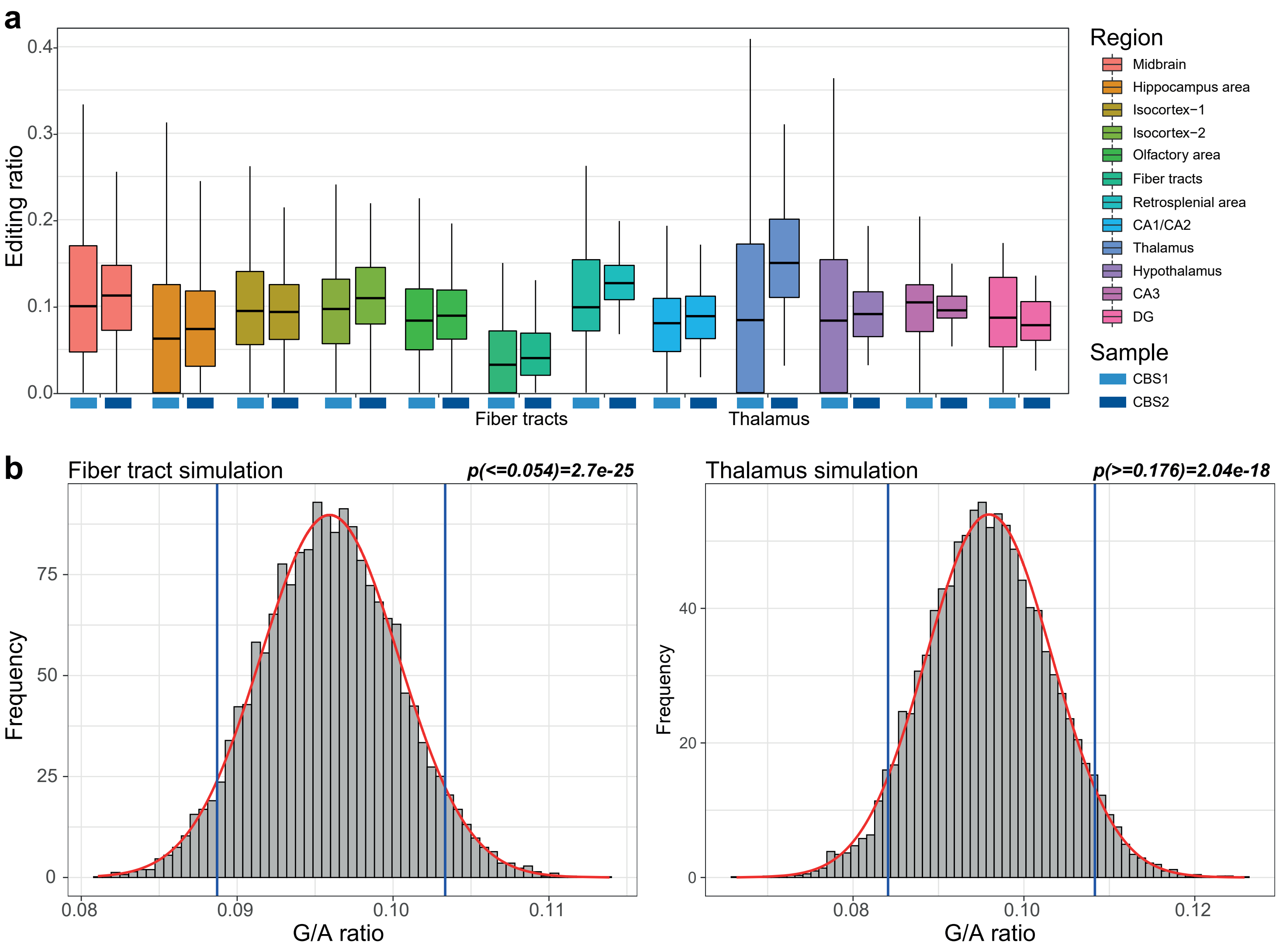


**Supplementary Figure 12. Resampling of global editing ratios and test statistics.** (**a**) Editing ratios were computed for each spatial region (CBS1: left boxplots, CBS2: right boxplots). (**b**) All capture spots across the CBS2 sample were resampled by randomly permuting the region labels 10,000 times. A normal distribution was fitted to the simulated editing ratios to calculate the probability of observing a value equal to, or more extreme, than the observed value.
