## Supplementary Figure 13. Correlation of editing ratios between coronal brain sections CBS1 and CBS2 (a) and between short and long-reads (b) for "The spatial landscape of gene expression isoforms in tissue sections"

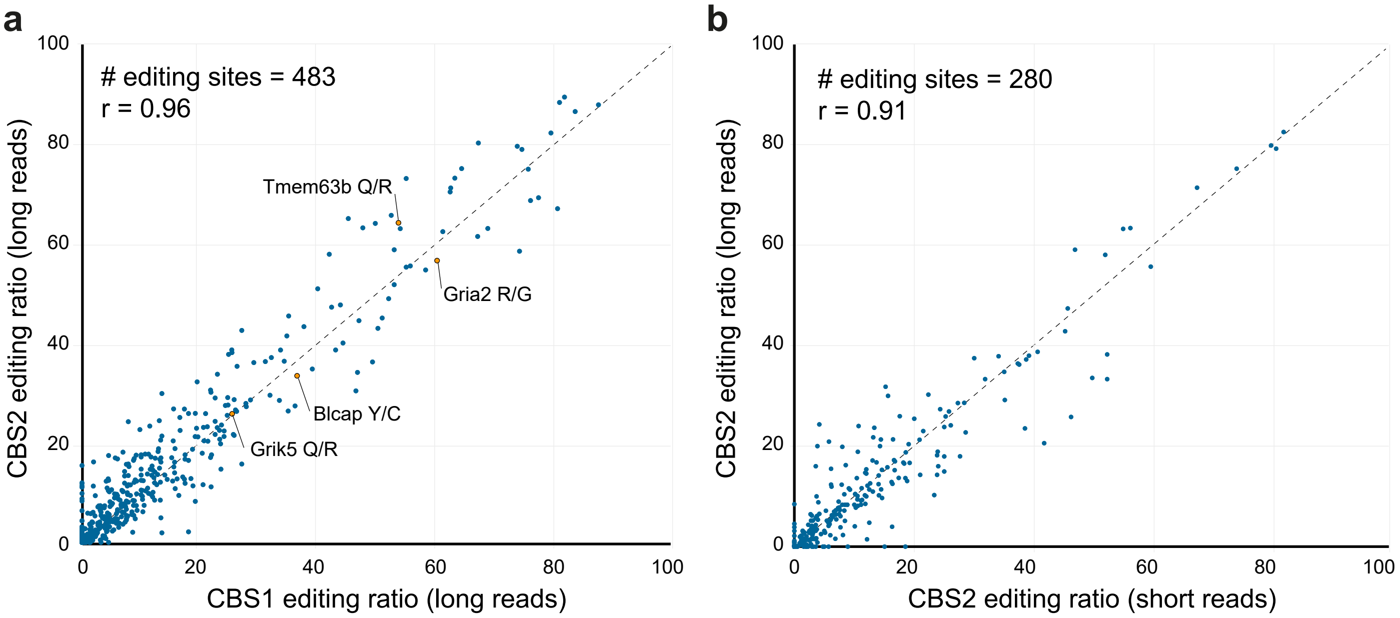


**Supplementary Figure 13. Correlation of editing ratios between coronal brain sections CBS1 and CBS2.** Scatter plots showing the editing ratios for editing sites backed by at least 20 UMIs in CBS1 and CBS2 for (**a**) Nanopore long read and (**b**) Illumina short read dataset.
