## Supplementary Figure 14. Coronal Brain Section (CBS2) spatial spots classification after SCTransform normalization for "The spatial landscape of gene expression isoforms in tissue sections"

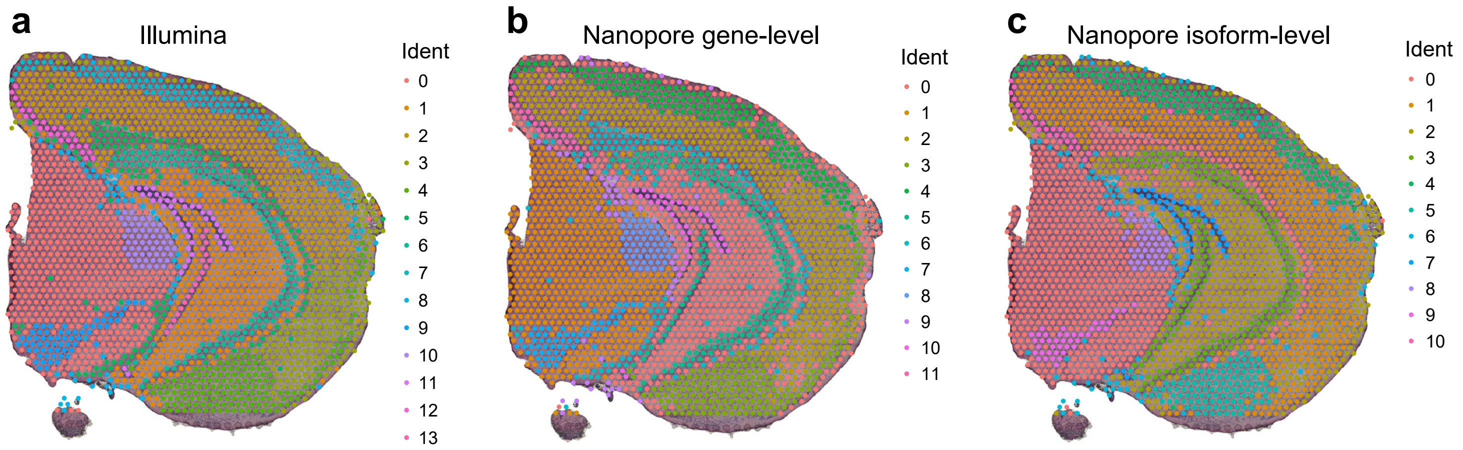


**Supplementary Figure 14. Coronal Brain Section (CBS2) spatial spots classification after SCTransform normalization.** The first 30 principal components of PCA were used for UMAP representation and clustering (resolution=0.4) using (**a**) Illumina “Spatial“ assay; (**b**) Nanopore gene-level “ISOG“ assay; and (**c**) Nanopore isoform-level “ISO“ assay.
